# Non-gonadal PIWI protein, Aubergine, regulates regenerative stem cell proliferation and tumorigenesis in the *Drosophila* adult intestine

**DOI:** 10.1101/2024.11.04.621896

**Authors:** Karen Bellec, Lynsey R Carroll, Kathryn AF Pennel, Yuanliangzi Tian, Yachuan Yu, Aslihan Bastem Akan, Caroline V Billard, Nora Doleschall, Alexander R Cameron, Fabiana Herédia, Alisson M Gontijo, Anna M Ochocka-Fox, James P Blackmur, Farhat V N Din, Malcolm G Dunlop, Joanne Edwards, Kevin Myant, Rippei Hayashi, Julia B Cordero

## Abstract

The PIWI-interacting RNA (piRNA) biosynthesis pathway is best studied for its role suppressing *Drosophila* germline transposable elements. Piwi, the founding member of the pathway, is involved in adult intestinal stem cell (ISC) homeostasis. Whether a broader role of the PIWI pathway exists in the intestine, remains unknown. Here, we characterise a role of the PIWI family protein Aubergine (Aub) in ISCs. While dispensable for basal ISC self-renewal, upregulation of Aub by damage-induced reactive oxygen species drives regenerative ISC proliferation through increased protein synthesis, including translation of ISC factors Myc and Sox21a. Unexpectedly, such roles of Aub in ISCs appear uncoupled from its piRNA regulatory function. Additionally, Aub and mammalian PIWIL1, mediate tumorigenic intestinal growth in *Drosophila* and human organoids, respectively. Our results reveal regulated protein translation as a fundamental aspect of regenerative ISC function and discover a central role of Aub in such process.

## Introduction

Since their discovery in *Drosophila melanogaster,*^1-3^ the PIWI protein family has been extensively studied for their conserved role protecting germline genome integrity via piRNA-dependent transposable elements (TEs) silencing.^4-13^ piRNAs are PIWI protein-bound 24-32 nucleotides long small non-coding RNAs derived from transposon-dense genomic loci called piRNA clusters^6^. PIWI proteins loaded with a piRNA exert gene silencing by cleaving mRNAs or recruiting chromatin modification enzymes to install heterochromatin. In the *Drosophila* germline, PIWI proteins Piwi and Aub mainly bind piRNAs that target TEs in the nucleus and the cytoplasm, respectively,^6,14-16^ while Argonaute 3 (AGO3) predominantly binds TE sense piRNAs to aid the production of TE antisense piRNAs through the ping-pong pathway.^6,7,17-20^

Numerous reports have documented the presence of PIWI proteins in somatic tissues and/or roles for these proteins beyond piRNAs and TE regulation^21-30,31^, including roles of Piwi in the maintenance of adult *Drosophila* intestinal stem cells.^28 27^ However, it remains unclear whether there is a general role of the PIWI pathway and piRNAs in the adult intestine or if PIWI proteins play roles distinct from their canonical piRNA-dependent function.

Here, we discovered that Aub is upregulated within the stem/progenitor compartment of the adult midgut in response to oxidative stress and is required to regulate regenerative and hyperplastic ISC proliferation. Despite the presence of piRNA-like molcules in ISCs/EBs our studies indicate that the induction of ISC proliferation by Aub is uncoupled from its canonical piRNAs regulatory function. Mechanistically, Aub promotes protein synthesis in regenerating ISCs, including translation of stem cell factors Myc and Sox21a. Furthermore, Aub or its mammalian orthologue PIWIL1, drive tumorigenic intestinal growth in the midgut and human intestinal organoids, respectively. Altogether, our results uncover non-canonical roles of Aub and PIWIL1 in physiological and pathological proliferation of the adult intestine in vivo.

## Results

### Aub is required for regenerative proliferation of ISCs following damage to the adult midgut epithelium

The adult *Drosophila* midgut epithelium is maintained and repaired by intestinal stem cells (ISCs). ^32,33^ Undifferentiated stem cell progeny, namely, enteroblasts (EBs) (**Fig1A, A’**) and pre-enteroendocrine cells are precursors of absorptive enterocytes (ECs) and secretory enteroendocrine cells (EEs) (**Fig1A’,A’’**), respectively.^34-36^ Visceral muscle (VM), terminal tracheal cells (TTCs) and enteric neurons (ENs) (**Fig1A’**) compose the intestinal microenvironment.^37,38^

RT-qPCR analysis of midguts from mated females showed low but detectable levels of *piwi*, *aub* and *ago3* mRNAs in homeostatic midguts, including the posterior midgut (R4-R5) (**Fig.1A, B**).^39,40^ We next assessed the role of the PIWI pathway in intestinal regeneration upon oral infection with the pathogen *Pseudomonas entomophila* (*Pe*)^41^ and quantified proliferating ISCs by phosphorylated histone H3 (PH3) staining (**Fig.1C** and **Fig.S1A**). *Pe* feeding caused robust ISC proliferation in wild type (*w^1118^*) flies when compared to their sucrose fed counterparts (**Fig.1C** and **Fig.S1A**). While loss of *piwi* mildly diminished intestinal regeneration (**Fig.1C** and **Fig.S1A**), *aub^HN2^/aub^QC42^*loss of function mutants^42,19,20,43^ depicted strong impairment in midgut regeneration (**Fig.1C** and **Fig.S1A**). Knockdown of *aub* within ISCs and EBs (stem/progenitors) by RNAi overexpression under the control of the temperature-sensitive *escargot-Gal4* (*esg^ts^*) abolished regenerative ISC proliferation (**Fig.1D** and **Fig.S1B**). On the other hand, ISC lineage tracing by mosaic analysis with a repressible cell marker (MARCM),^44^ revealed that, unlike *piwi,*^28^ knocking down *aub* did not cause any detectable effects in homeostatic ISC self-renewal (**Fig.S1C, D**). Unexpectedly, loss of *ago3*, an obligate partner of *aub* in the ping-pong pathway, had no impact on intestinal regeneration upon damage (**Fig.1C** and **Fig.S1A**). Similarly, we detected no effect on intestinal regeneration upon loss *spnE* (**Fig.S1E**), an essential ping-pong pathway RNA helicase.^20,45^ These data point to a distinctive role of *aub* in regenerative ISC proliferation of the adult *Drosophila* midgut.

**Figure 1:**
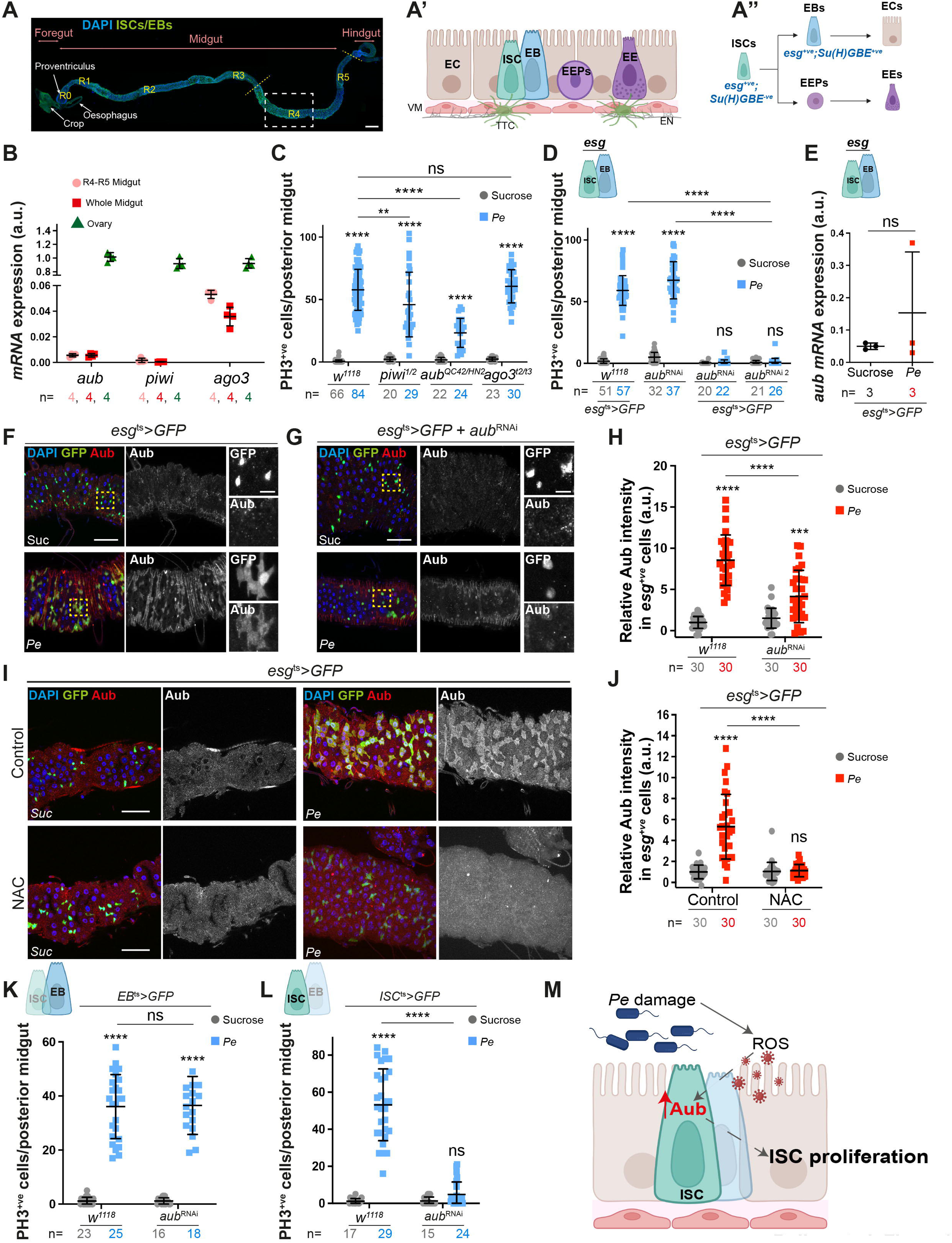
ISC Aub drives regenerative proliferation in the *Drosophila* midgut. **(A)** Mated female gut. ISCs and EBs (GFP; green). DAPI (blue) stains all nuclei. Yellow lines delineate posterior midgut, and the rectangle defines region of interest throughout the study. Scale bar = 200µm. **(A’, A”)** Schematics of midgut epithelium and associated tissues (A’); ISC lineage (A”). ISCs, intestinal stem cells; EBs, enteroblasts; EEPs, enteroendocrine cell precursors; ECs, enterocytes; EEs, enteroendocrine cells; VM, visceral muscle; EN, enteric neurons; TTC, terminal tracheal cell. **(B)** *aub*, *piwi* and *ago3* expression in midgut and ovary. n= biological replicates. **(C)** PH3 cells in midguts from *w^1118^, aub*, *piwi* or *ago3* mutant flies. **(D)** PH3 cells in control midguts (*esg^ts^>GFP* or UAS-*aub^RNAi^*only) or overexpressing independent *aub-RNAis* within ISCs/EBs (*esg^ts^>GFP + aub^RNAi^* or *aub^RNAi2^*). **(E)** *aub* expression in sorted ISCs/EBs. Mann-Whitney t test. n= biological replicates. **(F, G)** Aub staining (red and grey) in control or *aub^RNAi^* midguts. Yellow squares delineate the magnified area in righthand panels. **(H)** Quantification of data in F, G. n = number of cells. **(I)** Aub staining (red; grey) in midguts upon Sucrose or *Pe* feeding, with or without the antioxidant *N-acetyl cysteine* (NAC). **(J)** Quantification of data in I. n = number of cells. **(K, L)** PH3 cells in midguts overexpressing *aub^RNAi^* in either EBs (K) or ISCs (L). **(M)** Schematic of Aub regulation and function in regenerating ISCs. Unless otherwise noted, two-way ANOVA followed by Sidak’s multiple comparisons tests were used for statistical analysis and n = number of midguts/flies. a.u.; arbitrary units. Data are represented as mean +/- SD. ns, not significant; \*\**P*□*<*□0.01; \*\*\*\**P*□*<*□0.0001. Scale bars = 50µm.

### Damage-induced oxidative stress drives post-transcriptional upregulation of Aub in regenerating stem/progenitor cells of the adult *Drosophila* midgut

RT-qPCR experiments in sorted ISCs/EBs, identified by their enrichment of *esg* mRNA expression (**Fig.S1F**), revealed overall enrichment of *aub* in sorted cells versus whole midgut values (**Fig.1B, E**), but it did not show significant changes in *aub* mRNA upon *Pe* infection (**Fig.1E**). On the other hand, protein immunostaining experiments revealed strong upregulation of Aub expression in ISCs/EBs upon midgut damage (**Fig.1F, H**), which was markedly reduced following *esg-Gal4* driven RNAi *aub* knockdown (**Fig.1G, H**).

Intestinal damage triggered by pathogenic bacterial infection generates high levels of reactive oxygen species (ROS) in the gut lumen, mainly via enterocytes (ECs), as a protective host mechanism against the pathogen.^46^ ROS also play an important role as a signalling molecule influencing the production, secretion and stability of intestinal and niche derived factors necessary to induce ISC proliferation during midgut regeneration.^47,48^ Consistently, blocking ROS in *Pe* infected midguts by feeding animals with the antioxidant *N-acetyl cysteine* (NAC) led to significant impairment of Aub upregulation upon damage (**Fig.1I, J**). However, blocking ISC proliferation in the precence of infection did not impact Aub upregulation (**Fig. S1G-J**). These results suggest that pathogenic damage-induced upregulation of Aub is dependent on oxidative stress and precedes the activation of regenerative ISC proliferation.

### Aubergine works cell-autonomously in ISCs to drive midgut regeneration upon damage

Similarly to mammalian Paneth cells, EBs are components of the *Drosophila* intestinal stem cell niche.^49,50^ Wg/Wnt secretion from EBs is essential to paracrinally induce regenerative ISC proliferation.^50^ To distinguish the cell type where *aub* functions to regulate ISC proliferation, we induced gene knockdown in EBs or ISCs only using cell specific drivers *Su(H)GBE-Gal4,UAS-GFP; tub-Gal80*^ts^ or *esg-Gal4,UAS-GFP; Su(H)GBE-Gal80, tub-Gal80*^ts^, respectively (hereafter referred to as *EB^ts^* and *ISC^ts^*, respectively; **Fig.1K, L**).^51,52^ While no impact on intestinal regeneration was observed upon knocking down *aub* in EBs (**Fig.1K**), gene knockdown in ISCs was sufficient to recapitulate the impairment in regeneration observed upon dual ISCs/EBs *aub* knockdown (**Fig.1L**). Collectively, these results suggest that the role of Aub in midgut regeneration is ISC autonomous (**Fig.1M**). Consistently, Aub knockdown did not affect the production of the EB-derived Wg (Fig.S1K-M).

### Midgut stem/progenitor cells express TE mapping piRNA-like small RNAs, which are not affected upon Aub knockdown

The fact that other key components of the piRNA amplification/ping-pong pathway did not mimic the observed *aub* phenotype (**Fig.1C** and **Fig.S1A**) led us to hypothesize that Aub may be working in a non-canonical fashion to induce regenerative ISC proliferation in the adult midgut. Next, we took multiple complementary approaches to understand the mechanisms of action of Aub in the adult *Drosophila* midgut.

The presence of 2′-*O*-methylation at the 3′ ends of piRNAs confers resistance to sodium periodate oxidation.^9,53-55^ Consequently, RNA samples can be enriched with siRNAs and piRNAs versus other, and potentially more abundantly present, small RNAs populations. Although much less prominent than in oxidized libraries from ovaries, oxidation of whole gut libraries revealed an enrichment of 23-28 nucleotides long TE antisense small RNAs (**Fig.S2A-C**). Importantly, those RNAs showed nucleotide biases characteristic of piRNAs: Uridines at the 1^st^ base position of TE antisense reads (AS) and, to a lesser extent, Adenines at the 10^th^ base position of TE sense reads (S)—hallmarks of ping-pong piRNA biogenesis.^6^ Notably, this enrichment was not visible in previously published unoxidized midgut libraries,^56 28^(**Fig.S2D**) emphasising the importance of library oxidation to observe small enrichments in piRNAs. Stem/progenitor cells are underrepresented in bulk midgut tissue preparations, which include heterogenous cellular sub-types from the intestinal epithelium and associated microenvironment **(Fig. 1A’)**. We therefore next performed small RNA sequencing of oxidized samples from sorted ISCs/EBs (**Fig.2A**). Compared to whole midgut samples, small RNA sequencing from ISCs/EBs of control and regenerating midguts consistently showed a more pronounced piRNA signature (**Fig.2B-C’** versus **Fig.S2A, B**). However, this was not reduced by cell specific knockdown of *aub* (**Fig.2B-E**).

**Figure 2:**
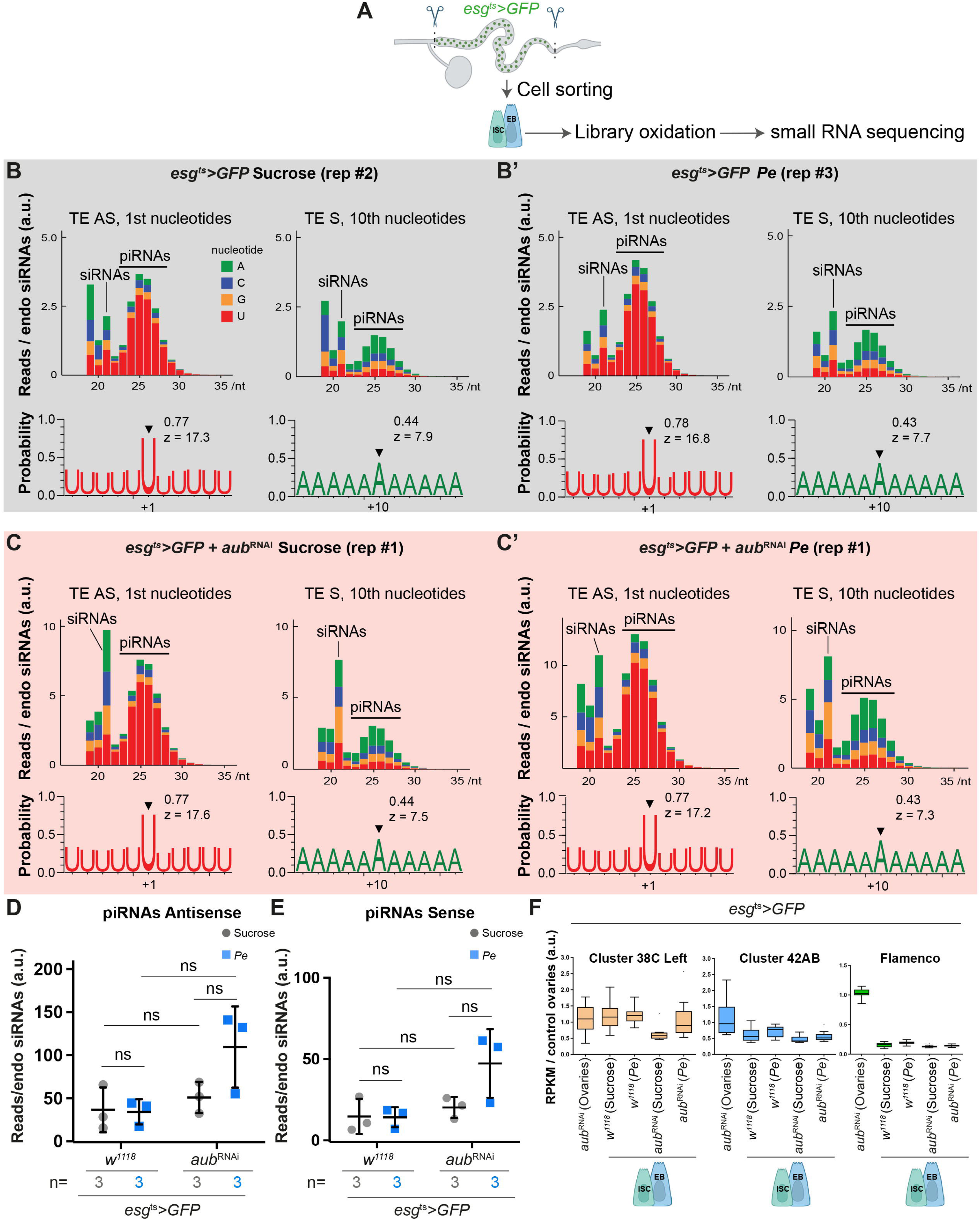
Aubergine does not impact the presence of piRNAs in *Drosophila* intestinal stem/progenitor cells. **(A)** Schematic of sorted ISCs/EBs (green dots) used for small RNA sequencing. **(B-C’)** Size distribution (top) of all transposon-mapping reads and Uridine- and Adenine-frequencies of piRNA size (>22nt) (bottom) transposon-mapping reads in oxidized libraries from sorted ISCs/EBs. **(D, E)** Quantification of TE antisense (D) and sense (E) piRNAs abundance as in B-C’. Two-way ANOVA followed by Sidak’s multiple comparisons tests were applied. n = 3 biological replicates. **(F)** Representation of piRNA clusters expression in sorted ISCs/EBs as in B-C’, compared to ovaries. Results are presented relative to control ovaries. a.u., arbitrary units. TE, transposable elements; AS, antisense; S, sense. ns, not significant. Data are represented as mean +/- SD.

Our analysis shows that as in ovaries, around 90% of all piRNA-like reads in ISCs/EBs mapped to TE sequences (**Fig.S2E**). We also find a strong bias (around 50%) towards ‘U’ at the immediate downstream nucleotide position of TE and non-TE mapping reads for all libraries, indicative of phased piRNA production (**Fig.S2F-H**).^18^ Despite these commonalities, ISCs/EBs appear to express piRNA-like populations, distinct from those expressed in the ovaries. The tiles analysis of genome-unique piRNA mappers in ISCs/EBs versus those in ovaries showed that piRNA germline-expressed clusters 38C and 42AB are comparatively more abundant in ISCs/EBs than those from the somatic cluster *flamenco* (**Fig.2F**). This is consistent across all conditions and genotypes analysed (**Fig.S3A-E**). In summary, our oxidised libraries from sorted adult midgut ISCs/EBs detected small RNAs with a molecular signature consistent with that of piRNAs. However, while effective to block regenerative ISC proliferation, RNAi dependent *aub* knockdown is not sufficient to deplete ISC/EB piRNA-like small RNAs.

### Aub regulates intestinal regeneration independently of its canonical piRNA regulatory function

We next performed mRNA sequencing from sorted ISCs/EBs (**Fig.S3F**) to measure the abundance of TE mRNAs in homeostasis and regeneration (**Fig.S3G, H**). While several TEs, such as *copia*, *roo* and *Doc*, were abundantly expressed in ISCs/EBs in both conditions, this was independent of Aub (**Fig.S3G, H**). These TEs are neither abundantly expressed nor under the control of Aub in the ovaries (**Fig.S3I**).^57^ Interestingly, we observed that *flea* expression, a TE regulated by Aub in the germline,^58^ (**Fig.S3I**) was upregulated in ISCs/EBs upon *Pe* infection regardless of Aub presence (**Fig.S3G, H**). These data confirms the distinctive nature of midgut versus ovary TEs and are consistent with recent work suggesting that stress-associated TE regulation in the midgut may be carried out by mechanisms independent of the piRNA machinery.^56^

Next, we used site-directed transgenesis to generate *UAS-aub^WT^*, *UAS-aub^AA^* or *UAS-aub^ADH^*lines (**Fig.S4A**). *UAS-aub^AA^* contains a double point mutation in the PAZ domain, responsible for the loading of piRNAs,^59^ while *UAS-aub^ADH^* carries a single mutation in the PIWI domain, responsible for Aub’s endonuclease activity and piRNA biogenesis.^19^ Mutations in *aub* have been previously correlated with female sterility, a reduced size of the ovaries and embryonic lethality.^19,42,59,60^ Consistent with previous reports,^19^ we observed that only *UAS-aub^WT^* could partially restore fertility when expressed in the germline (**Fig.S4B**). On the other hand, comparable levels of transgene expression in midgut ISCs/EBs (**Fig.S4C-G**) were similarly capable of improving regenerative ISC proliferation in *aub^HN2/QC42^*midguts (**Fig.3A**) and resulted in comparable gain of function phenotypes in undamaged, wild type midguts (**Fig.3B, C**). Our results suggest that Aub is necessary and sufficient to drive ISC proliferation in the adult *Drosophila* midgut and it does so independently of its canonical piRNA regulatory function.

**Figure 3:**
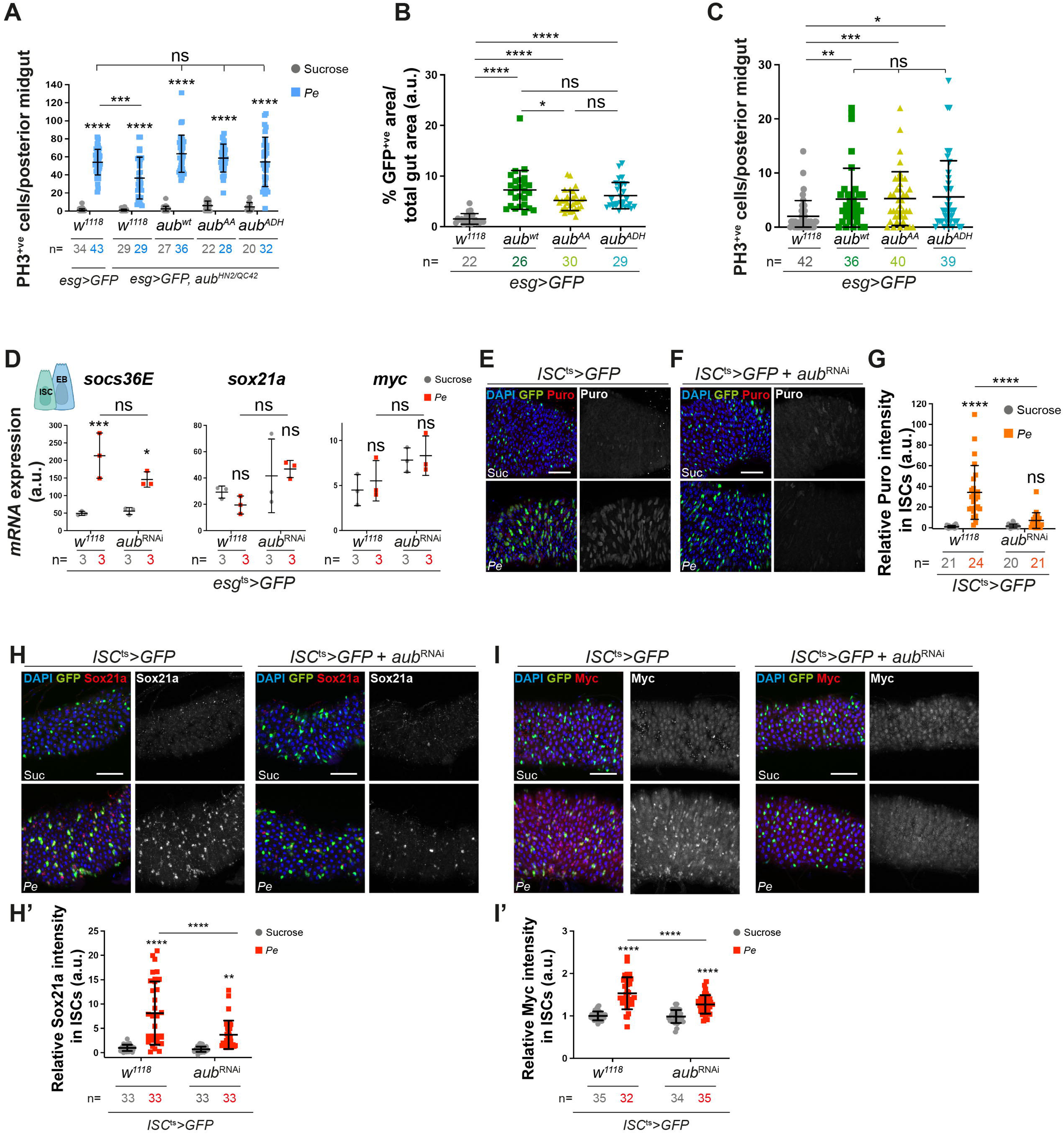
Aub regulates intestinal regeneration independently of its piRNA regulatory function. **(A)** PH3-positive cells in *aub^HN2^/aub^QC42^* midguts expressing GFP alone or with *aub^WT^*, *aub^AA^* or *aub^ADH^* within ISCs/EBs. **(B)** *esg>GFP* area in control ISCs/EBs or overexpressing *aub^WT^*, *aub^AA^*or *aub^ADH^*. Shapiro-Wilk normality test followed by Mann-Whitney t test. **(C)** PH3 cells in midguts as in B. Shapiro-Wilk normality test followed by a Kruskal-Wallis one-way ANOVA and Dunn’s multiple comparisons test. **(D)** Relative mRNA expression of *socs36e, sox21a and myc* in sorted ISCs/EBs. n = 3 biological replicates. **(E, F)** Puromycin staining (red and grey) in control ISCs (green) or expressing *aub^RNAi^*. **(G)** Quantification of data in E, F. **(H, I)** Sox21a (H) and Myc (I) staining (red and grey) in control ISCs (green) or expressing *aub^RNAi^*. **(H’, I’)** Quantification of data in H, I. Unless otherwise noted, two-way ANOVA followed by Sidak’s multiple comparisons tests were applied. n = number of midguts/flies quantified. a.u., arbitrary units. ns, not significant; \**P*□<□0.05, \*\**P*□*<*□0.01, \*\*\**P*□<□0.001 and \*\*\*\**P*□*<*□0.0001. Data are represented as mean +/- SD. Scale bars= 50µm.

### Aub regulates ISC proliferation through the translation of core regenerative factors in the adult midgut

To further characterize the processes underpinning the role of Aub in the midgut, we assessed its effect on regenerative signalling pathways. The EGFR/MAPK, Wnt and JAK-STAT pathways drive intestinal regeneration through ISC activation of their respective effectors, Sox21a, Myc and Socs36E.^41,49,50,61-66^ We hypothesized that these transcription factors would represent good candidates to mediate the cell autonomous function of Aub within ISCs (**Fig.1K-M**). Consistent with prior reports**,^41^** our RT-qPCR analysis showed upregulation of the transcriptional target of JAK/Stat signalling, *socs36e*, in sorted ISCs/EBs from *Pe* treated midguts, confirming the regenerative status of our sorted cell population (**Fig.3D**). However, we observed no significant impact of *aub* knockdown on *socs36e* mRNA. In contrast*, sox21a* and *myc* mRNA expression in sorted cells did not change upon *Pe* infection and was overall increased—albeit not statistically significantly—in Aub-depleted ISCs/EBs (**Fig.3D**). Neither of these scenarios explained Aub-dependent ISCs proliferation upon damage.

Biological functions of Aub in the germline and embryo include control of the translation of mRNAs coding for specific factors.^43,67-70^ Puromycin incorporation experiments to measure global protein translation,^71-73^ showed a significant increase in protein synthesis in regenerating midguts, which was highly dependent on *aub* expression (**Fig.3E-G**). Consistent with prior reports ^50,74^, we observed significant Sox21a (**Fig.3H, H’**) and Myc (**Fig.3I, I’**) protein upregulation in ISCs during midgut regeneration. Critically, this was significantly impaired upon *aub* knockdown (**Fig.3H-I’**).

The preservation of proteostasis, the balance between protein synthesis and degradation, is an important aspect of intestinal homeostasis in the *Drosophila* midgut.^75^ Levels of Sox21a and Myc are dynamically regulated during damage-induced ISC proliferation and post-damage recovery,^47^ with a significant decay of both proteins observed 24hs after removal of the damaging agent (**Fig.S5A, C**). However, overexpression of Aub did not prevent Sox21a or Myc downregulation after damage (**Fig.S5B, D**).

Knocking down the *proteasome* β*5* subunit (*pros*β*5)*,^76^ in ISCs/EBs to inhibit protein degradation, leads to the accumulation of multi-ubiquitinated proteins in damaged midguts (**Fig.S5E, F**). We reasoned that an increase in protein degradation upon *aub* knockdown would become evident by blocking the proteasomal machinery. However, we observed no significant difference in the levels of accumulated multi-ubiquitinated upon *aub* knockdown (**Fig.S5E, F**), suggesting that Aub is not a major regulator of protein stability within ISCs during midgut regeneration.

We next assessed the capacity of *UAS-aub^WT^*, *UAS-aub^AA^*or *UAS-aub^ADH^* to regulate Sox21a and Myc expression in the midgut. Overexpression of either isoform of Aub was similarly sufficient to induce mild levels of Myc and Sox21a in homeostatic wild type midguts (**Fig.4A-C**) and to significantly restore protein upregulation in *aub^HN2/QC42^* midguts following pathogen-induced damage (**Fig.4D-F**). Altogether, these data suggest that Aub drives regenerative ISC proliferation through cell autonomous upregulation of protein synthesis, including inducing *sox21a* and *myc* translation, either directly or indirectly and in a manner that is independent of its association with piRNAs (**Fig.4G**).

**Figure 4:**
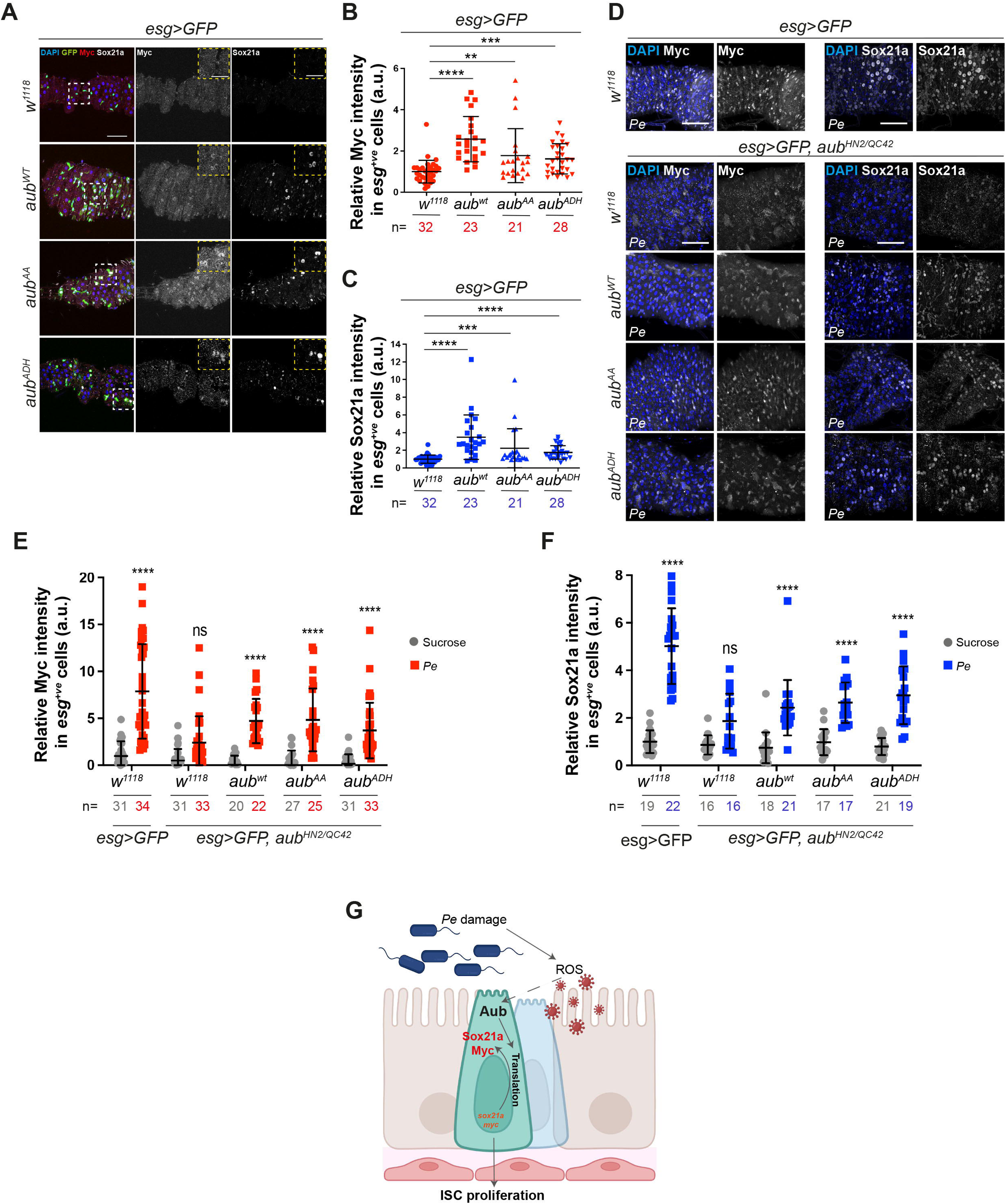
Aub regulates core regenerative stem cell factors in the adult midgut. **(A)** Myc and Sox21a staining in ISCs/EBs from control flies or flies overexpressing *aub^WT^*, *aub^AA^*or *aub^ADH^*. Dashed white squares delineate magnified areas shown in the dashed yellow panels. **(B, C)** Quantification of Myc (B) and Sox21a (C) staining as in A. Shapiro-Wilk normality test followed by Mann-Whitney t test. **(D)** Myc and Sox21a staining in ISCs/EBs from *aub^HN2^/aub^QC42^* midguts expressing *aub^WT^*, *aub^AA^* or *aub^ADH^* in ISCs/EBs. **(E, F)** Quantification of staining as in (D). Two-way ANOVA followed by Sidak’s multiple comparisons test. n = number of midguts/flies. a.u., arbitrary units. **(G)** Schematic of Aub regulation and its function on Sox21a and Myc induction in regenerating ISCs. Data are represented as mean +/- SD. ns, not significant (*P*□>□0.05); \**P*□<□0.05, \*\**P*□*<*□0.01, \*\*\**P*□<□0.001; \*\*\*\**P*□*<*□0.0001. Scale bars=50µm.

### Aub regulates ISC proliferation through selective interaction with subunits of the translation initiation machinery

Previous studies showed that Aub promotes translation of key developmetal genes in the germline by interacting with translation initiation complexes eukaryotic initiation factor complex 3 (eIF3) and 4 (eIF4).^69,70^ Immunostaining revealed significant upregulation of eIF3C (**Fig.5A-B** and **Fig.S6A**) and eIF4G (**Fig.5C-D** and **Fig.S6B**) subunits in ISCs/EBs in response to damage.

**Figure 5:**
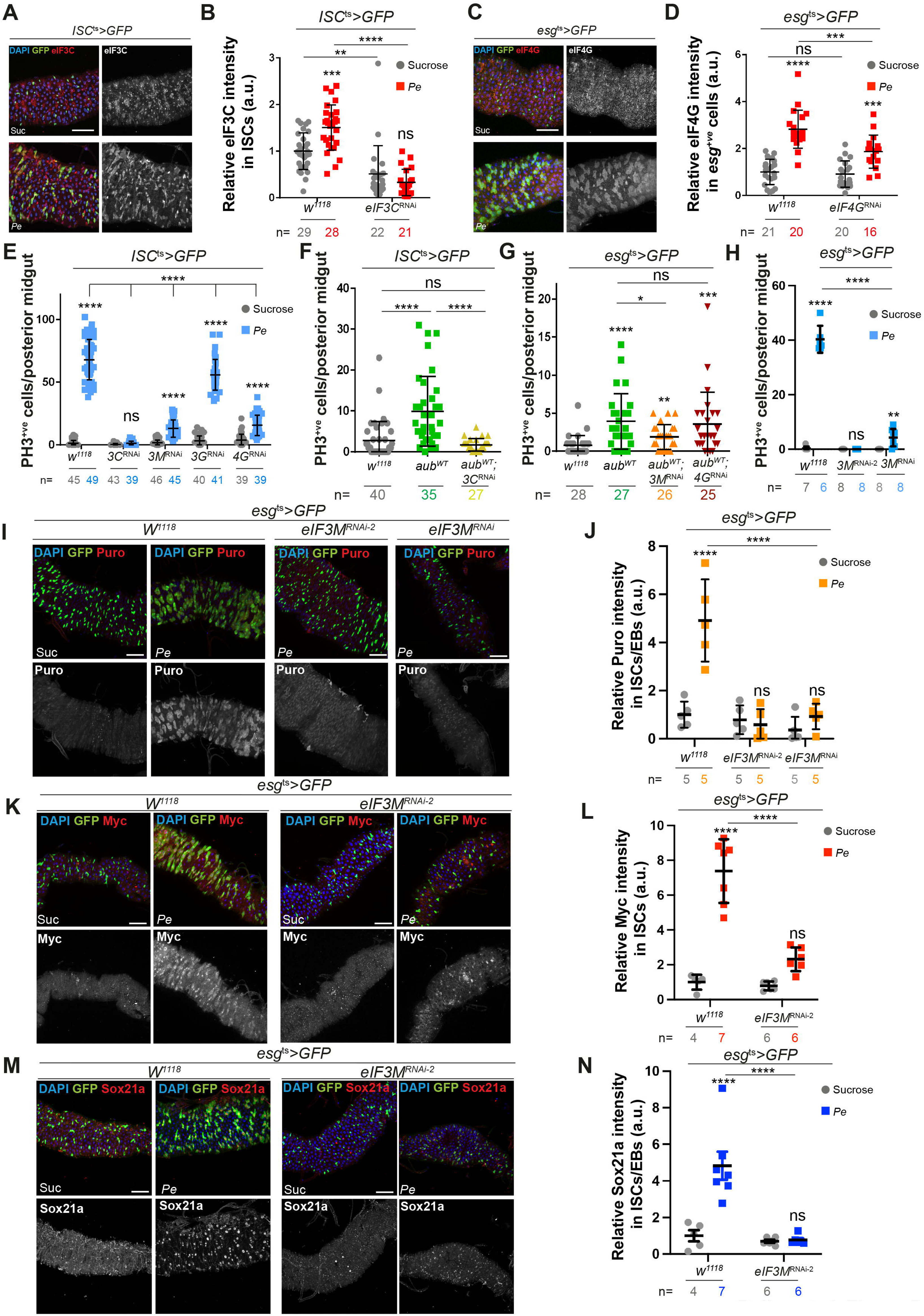
Aub interacts with selective eIF3 subunits to regulate ISC function during midgut regeneration. **(A)** eIF3C staining (red and grey) in ISCs (green). **(B)** Quantification of eIF3C staining as in A and upon *eIF3c* RNAi overexpression in ISCs. **(C)** eIF4G staining (red and grey) in ISCs/EBs (green). **(D)** Quantification of eIF4G staining in midguts as in C and upon *eIF4G* RNAi overexpression in ISCs/EBs. **(E)** PH3 in midguts expressing RNAi against eIF3 complex subunits or eIF4G within ISCs. **(F)** PH3-positive cells in midguts expressing *aub^WT^* or *aub^WT^* with *eIF3C^RNAi^* within ISCs. Shapiro-Wilk normality test followed by a Kruskal-Wallis one-way ANOVA and Dunn’s multiple comparisons test. **(G)** PH3-positive cells in midguts expressing *aub^WT^* or *aub^WT^* with *eIF3M^RNAi^* or *eIF4G^RNAi^* within ISCs/EBs. Shapiro-Wilk normality test followed by Mann-Whitney t test. **(H)** PH3 in midguts expressing independent *eIF3M^RNAi^* lines in ISCs/EBs. **(I)** Puromycin staining (red and grey) in ISCs/EBs (green) of midguts as in H. **(J)** Quantification of data as in I. **(K, M)** Myc (K) and Sox21a (M) staining (red and grey) in ISCs/EBs (green) expressing GFP or *eIF3M^RNAi-2^*. (**L, N**) Quantification of staining as in K and M, respectively. Unless otherwise noted, two-way ANOVA followed by Sidak’s multiple comparisons tests were applied. n = number of midguts/flies. a.u., arbitrary units. Data are represented as mean +/- SD. ns, not significant; \**P*□<□0.05, \*\**P*□*<*□0.01, \*\*\**P*□<□0.001; \*\*\*\**P*□*<*□0.0001. Scale bars are 50µm.

Next, we measured regenerative ISC proliferation upon RNAi dependent knockdown of *eIF3* and *eIF4* subunits *eIF3C, eIF3M, eIF3G* ^77-78^ and *eIF4G,* a previously known binding partner of Aub in the germline (**Fig.5E**).^69^ Experiments on *eIF3B, eIF3D, eIF4A* and *eIF4E* knockeddown did not progress due to high animal lethality or homeostatic ISC loss. Among the factors whose depletion did not affect ISCs homeostasis, RNAi against *eIF3C* and *eIF3M* impaired ISCs proliferation upon infection or Aub overexpression. In contrast, RNAi against *eIF3G* and *eIF4G* did not impact regenerative ISCs proliferation, or only affected damage-induced ISCs proliferation while having minimal impact on Aub-induced ISC proliferation (**Fig.5E-G**).

Consistent with its impact on regenerative (**Fig. 5E, H**) and Aub-induced ISCs proliferation **(Fig. 5G)**, knocking down *eIF3M* significantly impaired protein translation (**Fig. 5I, J**) and the upregulation of Sox21a and Myc following midgut damage (**Fig.5K-N** and **Fig. S6E-H**). Interestingly, while *eIF3c* knockdown efficiently inhibited protein translation (**Fig.S6C, D**), intestinal regeneration (**Fig.5E**) and Aub-induced ISCs proliferation (**Fig.5F**), it did not show detectable impact on damage-induced upregulation of either Sox21a or Myc in ISCs (**Fig.6A-D**). Suggesting that, unidentified targets of eIF3C, other than Sox21a and Myc, promote its role on regenerative and Aub-dependent ISCs proliferation. Further investigation into the relationship between Aub and eIF3C revealed that the observed upregulation of eIF3C upon midgut damage (**Fig.5A-B**) was post-transcriptional (**Fig.6E**) and dependent on Aub (**Fig.6F-H**). Notably, Aub knockdown did not affect eIF4G induction (**Fig. S6I, J**). Overall, these results indicate a functional link between Aub and eIF3C that does not globally extend to core translation initiation factors. Furthermore, they point to a selective rather than general role of Aub regulating protein translation in the regenerating midgut. Consistently, knocking down *aub* had no effect on the upregulation of Armadillo/β-Catenin (**Fig. S6K, L**), a conserved component of canonical Wnt signalling, that regulates gene transcription, including Myc, during intestinal regeneration and cancer ^79-80^. This suggests that Aub activate Myc expression downstream of Armadillo/β-Catenin.

**Figure 6:**
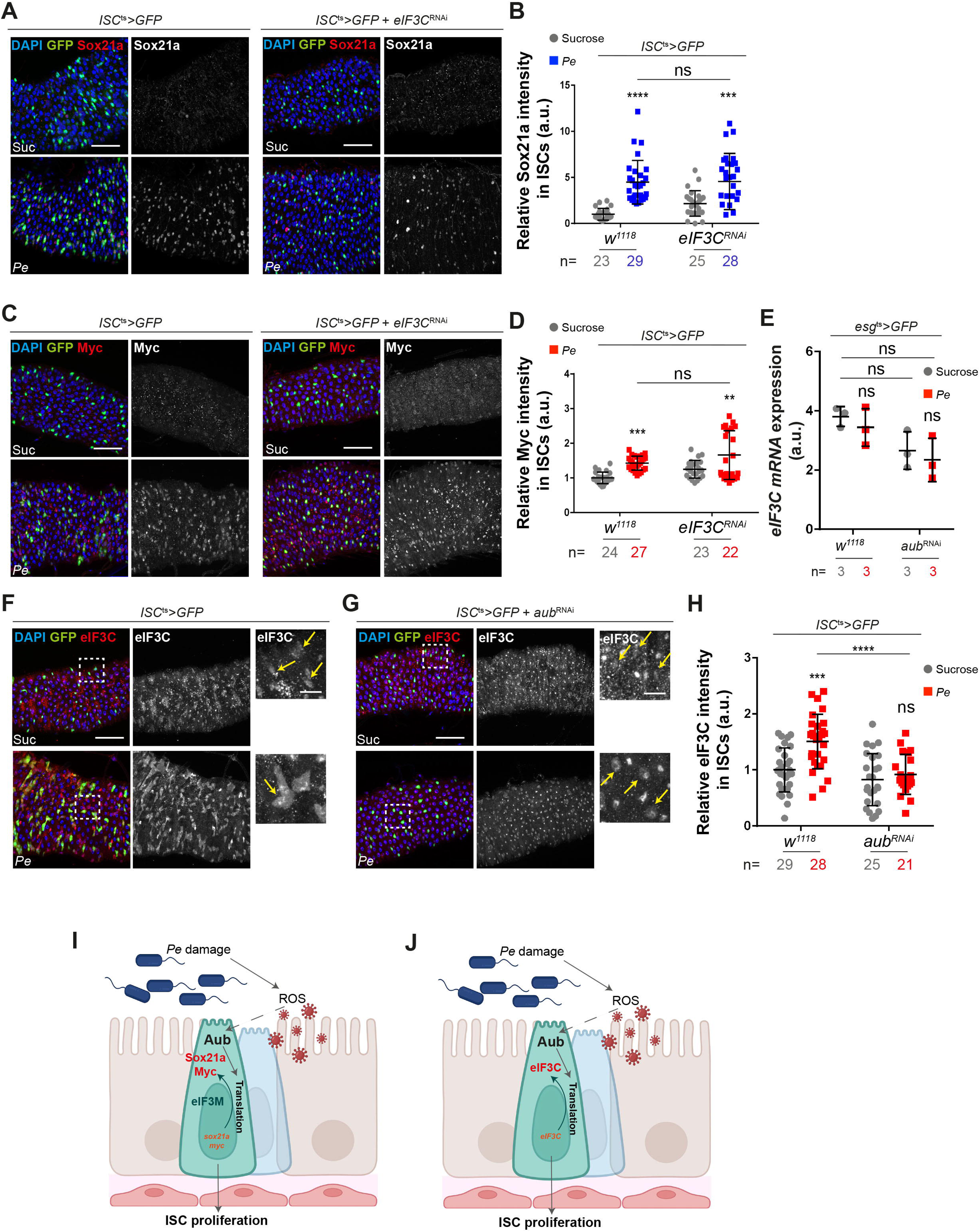
Aub interacts with selective eIF3 subunits to regulate ISC function during midgut regeneration. **(A)** Sox21a staining (red and grey) in ISCs expressing *eIF3C^RNAi^* (green). **(B)** Quantification of staining as in A. **(C)** Myc staining (red and grey) in ISCs expressing *eIF3C^RNAi^* (green). **(D)** Quantification of staining as in C. **(E)** Relative mRNA expression of *eIF3C* in control or *aub^RNAi^* sorted ISCs/EBs. n = 3 biological replicates. **(F, G)** eIF3C staining (red and grey) in ISCs expressing *aub^RNAi^* (green; yellow arrows). **(H)** Quantification of data as in F, G. **(I, J)** Schematic working models of proposed interrelationship between Aub and eIF3M (I) or eIF3C (J) during regenerative ISC proliferation. Unless otherwise noted, two-way ANOVA followed by Sidak’s multiple comparisons tests were applied. n = number of midguts/flies. a.u., arbitrary units. Data are represented as mean +/- SD. ns, not significant; \**P*□<□0.05, \*\**P*□*<*□0.01, \*\*\**P*□<□0.001; \*\*\*\**P*□*<*□0.0001. Scale bars= 50µm.

Altogether, these results suggest that Aub regulates regenerative ISC prolieration by inducing, directy or indirectly, the translation of a subset of mRNAs, including *sox21a*, *myc* and *eIF3C* (**Fig. 6I, J**). Furthermore, the role of Aub on ISCs involves functional interaction with specific subunits of the translation initiation complex.

### Aub and PIWIL1 drive ISC proliferation in CRC

Growing evidence in humans suggests misexpression of PIWI proteins and/or piRNAs in multiple cancers ^81-91^ including colorectal,^92-95^ and gastric cancers,^82,96^. We used a PIWIL1 antibody (**Fig.S7A, B**) to stained tissue samples from CRC patient samples, which showed strong expression of cytoplasmic PIWIL1 in tumour tissue from advanced stage cancers (**Fig.S7C**). Consistently, our analysis of *PIWIL1* gene expression in colon tissue samples revealed significantly higher *PIWIL1* gene expression in colon adenocarcinoma versus normal tissues (**Fig.7A, B**). Microarray data analysis revealed strong overexpression of *PIWIL1* in aggressive metastatic colon tumours (**Fig.7C**). Analysis of *PIWIL1* expression across different CRC subtypes,^97^ showed *PIWIL1 gene* expression significantly upregulated in the Consensus Molecular Subtype 1 (CMS1) human CRC subtype (**Fig.7D**), which presents high microsatellite instability correlated with hypermethylated profiles and strong immune activation.^97^ Consistently, we observed significant *PIWIL1* upregulation in patients with tumours characterised by high microsatellite instability (**Fig.7E**).

**Figure 7:**
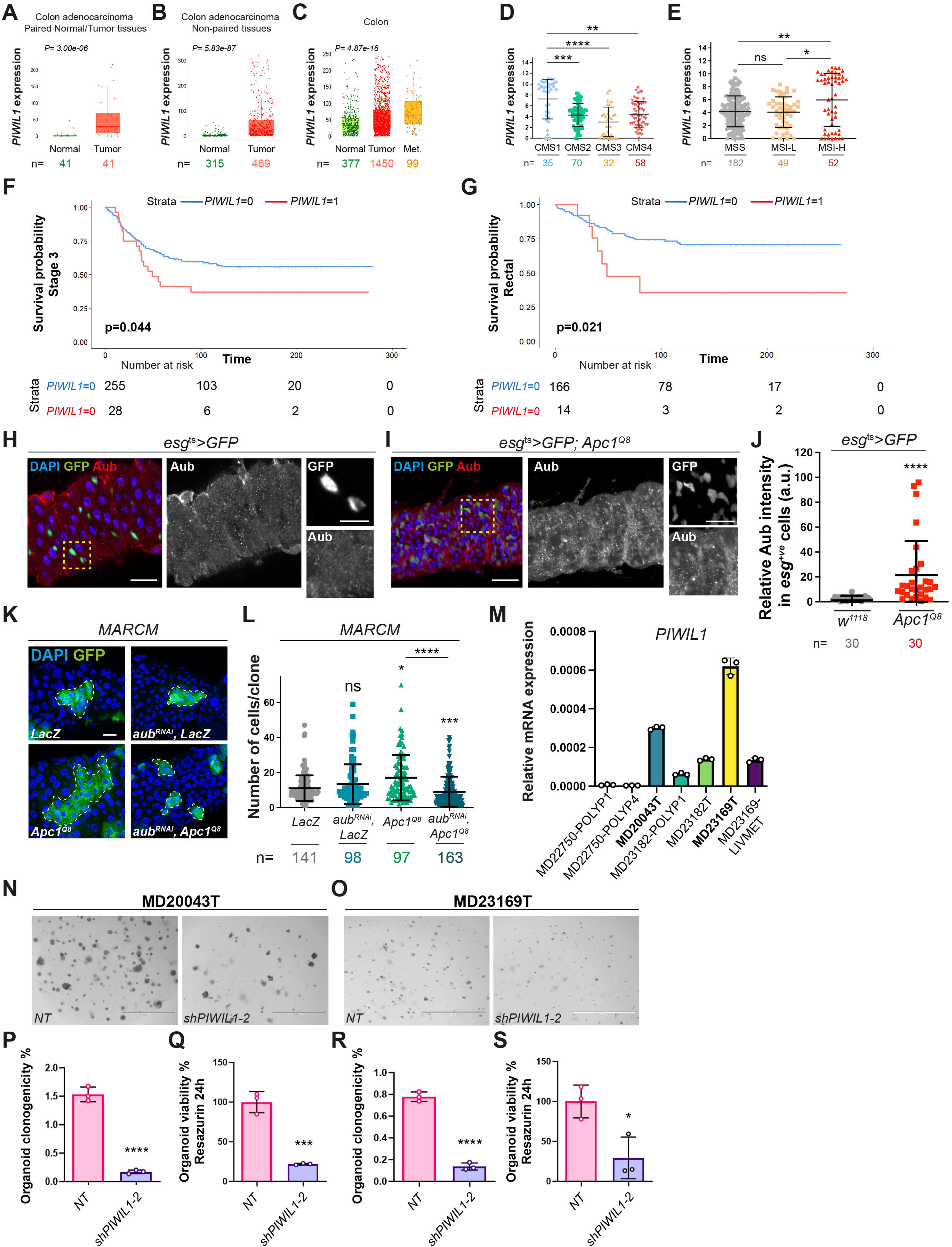
Aub and PIWIL1 mediates ISC proliferation in CRC. (A, **B)** *PIWIL1* expression in normal and tumour tissues from human colon adenocarcinoma. Mann-Whitney t test. n = number of patients. **(C)** *PIWIL1* expression in normal, tumour and metastatic tissues from human colon adenocarcinoma. Kruskal-Wallis one-way ANOVA test. n = number of patients. Box plots: line indicates median, and box represents the values included between the 25th and 75th percentiles. **(D)** *PIWIL1* expression across 4 CRC subtypes. Shapiro-Wilk normality test followed by a Kruskal-Wallis one-way ANOVA and Dunn’s multiple comparisons tests. n = number of patients. **(E)** *PIWIL1* expression according to microsatellite status of cancers (TCGA, PanCancer Atlas). MSS: microsatellite stability; MSI-L: low microsatellite instability; MSI-H: high microsatellite instability. Shapiro-Wilk normality test followed by a Kruskal-Wallis one-way ANOVA and Dunn’s multiple comparisons tests. n = number of patients. **(F, G)** Kaplan-Meier survival analysis of *PIWIL1* expression in a cohort of CRC patients (n=787) with stage 3 disease (F) and with tumours located within the rectum (G). **(H, I)** Aub staining (red) in ISCs/EBs (green) from control (I) or *Apc^1Q8^*mutant midguts (I). Dashed yellow squares delineate the high magnification shown in righthand panels. **(J)** Quantification of staining as in H, I. Shapiro-Wilk normality test followed by Mann-Whitney t test. n = number of cells quantified. **(K)** MARCM clones (green) expressing *LacZ* (control) or in combination with *aub^RNAi^*, and *Apc^1Q8^* clones with or without *aub^RNAi^*. Dashed white lines delineate clonal margins. Scale bars = 50µm **(L)** Number of cells per clone as in K. Shapiro-Wilk normality test followed by Kruskal-Wallis one-way ANOVA and a Dunn’s multiple comparisons tests. n = number of clones quantified. **(M)** Relative *PIWIL1* mRNA expression in human intestinal organoids derived from polyps or from colorectal tumours. n = 3 biological replicates. **(N, O)** Patient-derived CRC organoids transduced with a non-targeted control (NT) or *shPIWIL1-2* RNAi. Scale bars = 1000µm **(P-S)** Organoid clonogenicity upon NT or *shPIWIL1-2* RNAi transduction. Unpaired t tests. n = 3 biological replicates. a.u., arbitrary units. In all cases, data are represented as mean +/- SD. ns, not significant; \**P*□<□0.05, \*\**P*□*<*□0.01, \*\*\**P*□<□0.001; \*\*\*\**P*□*<*□0.0001. *LIVEMET*: Liver metastasis.

Survival analysis from a large local cohort of CRC patients demonstrated a significant association between high *PIWIL1* RNA expression and reduced cancer specific survival only in patients with stage 3 disease (**Fig.7F** and **Fig. S7D, E**), which was potentiated in patients with rectal tumours (**Fig.7G**). Altogether, our results confirm a positive correlation between *PIWIL1* and CRC and identify specific CRC subtypes where *PIWIL1* overexpression is most prominent, suggesting potential selectivity in the role of *PIWIL1* towards highly aggressive intestinal tumours.

We next looked at Aub expression in a CRC-like fly model driven by the loss of the tumour suppressor *Adenomatous polyposis Coli* (*Apc)*, a major driver of early intestinal tumourigenesis, whose loss leads to hyperactivation of Wnt signalling through constitute activation of β-catenin and upregulation of Myc.^80,98,99^ Aub expression was increased in ISCs/EBs from hyperplastic *Apc1* mutant midguts (**Fig.7H-J**).^80,100^ We next used the MARCM lineage tracing system ^44^ to induce ISC control (*LacZ*) or *Apc1* mutant clones upon *aub* knockdown (*aub^RNAi^*) (**Fig.7K, L**). While *aub* knockdown did not impact homeostatic clonal growth (**Fig.7K, L**), it strongly suppressed overgrowth of *Apc1* clones (**Fig.7K, L**) and ISC proliferation following overexpression of the Wnt ligand Wingless (Wg) ^50,66,80^ (**Fig.S7F**). Together with our regeneration data, these results re-enforce the essential role of ISC Aub mediating tumourigenesis downstream of Wnt ligand and β-catenin activity.

Screening a panel of human CRC organoids for *PIWIL1* expression revealed highest levels of gene expression in organoids derived from aggressive tumour types when compared to those obtained from benign polyps (**Fig.7M**). We used shRNA dependent gene knockdown to address the functional role of *PIWIL1* in the two patient-derived CRC organoid lines that showed highest gene expression levels (**Fig.7N, O**). Two independent shRNA constructs showed significant *PIWIL1* knockdown, with one shRNA (*shPIWIL1-2*) leading to at least 70% gene knockdown efficiency in organoid lines (**Fig.S7G, H**). Knocking down *PIWIL1* resulted in strong impairment of organoid clonogenicity/proliferative capacity (**Fig.7P, R**), and survival (**Fig.7Q, S**) without affecting organoid stemness as assessed by expression of the ISC marker *Lgr5* (**Fig.S7I, J**). In line with our *Drosophila* findings, these results suggest that PIWIL1 does not impair stem cell maintenance in the intestine, but it is required to drive ISC hyperproliferation and tumour growth in human CRC organoids. Altogether, our data suggest a stem cell intrinsic role of Aub inducing intestinal regeneration and tumorigenesis and a similar function of PIWIL1 driving human intestinal tumour growth. Importantly, we provide *in vivo* and *in vitro* paradigms amenable for broader mechanistic studies on the role of PIWI proteins and piRNAs in intestinal physiology and tumorigenesis.

## Discussion

Multiple lines of evidence have demonstrated roles of PIWI proteins beyond their well-defined function as repressors of TEs expression.^43,59,67-70,101-103^ Here, we present a non-canonical role of intestinal stem cell Aub, which involves the regulation of intestinal regeneration through translational control of conserved transcription factors and a core subunit of the translation initiation machinery.

### An inducible source of Aub regulates ISC proliferation in the regenerating adult *Drosophila* midgut

While levels of *aub* mRNA expression in the midgut are very low when compared to ovaries, pathogenic stress leads to striking upregulation of Aub protein in regenerating midguts (**Fig. 1**). Remarkably, this upregulation extends to exogenous overexpressed Aub coding sequences with minimal mRNA translation domains (**Fig.S4**), suggesting likely post-translational mechanism regulating Aub in the system. We identified ROS as a regulator of damage-induced Aub in the regenerating midgut (**Fig.1**). A conserved role of ROS on transcriptional and post-translational regulation of HIF1-17/Sima ^104, 47^ is associated with its influence in the remodelling of the vascular microenvironment of the regenerating midgut ^47^. As such, one possibility is that ROS may preserve Aub protein stability during intestinal repair.

### Aub acts as a regulator of protein translation in ISCs

Our work underscores a central role of inducible protein synthesis in the capacity of stem cells to trigger the regenerative response of the intestine following damage, as well as the complexity of the protein translation machinery operating within ISCs.

Results from our puromycin incorporation assay suggests that upregulation of protein translation is a prominent feature of the regenerative response of ISCs to damage and that Aub is required for a significant proportion of such response, including translation of key regeneration effector proteins, such as Sox21a and Myc. Questions emerging from our study include: Is damage-induced regeneration in general and Aub in particular linked to upregulation of global or mRNA specific translation? What are the mechanism of translation employed by regenerating ISCs?

Our data only partially addresses some of these questions. On the one hand, the striking inhibitory effect of Aub knockdown on puromycin incorporation and its necessity to upregulate the expression of the core translation initiation factor subunit eIF3C, may indicate a non-discriminatory role of Aub on global protein translation. However, we showed that knockdown of eIF3C it is not sufficient to impair translation of the two Aub targets *sox21a* and *myc* (**Fig. 6**). Furthermore, Aub knockdown does not impair the upregulation of two highly relevant drivers of regenerative ISC proliferation, eIF4G and Arm/β-Catenin (**Fig. S6**). This evidence suggest that, rather than global, the role of Aub on protein translation in regenerative ISCs is likely to involve the regulation of specific subsets of mRNAs.

While canonical mRNA translation involves assembly of the complete translation initiation machinery,^105^ differential requirement of translation initiation factors and target specific mRNA translation by the eIF3 and eIF4 have been reported in *Drosophila* and mice. ^106,107,77,108^ Consistently, our results show that eIF3C likely drives ISC proliferation upon damage through the regulation of a yet to be identified subset of mRNAs. On the other hand, the non-core eIF3M subunit induces broad protein translation, including Sox21a and Myc, in regenerating ISCs.

Physical interaction with target mRNAs or translation initiation factors subunits mediates the role of Aub in post-transcriptional gene regulation in the germline and ovaries.^73,77^ Unbiased molecular and biochemical approaches for global analysis of protein/protein and protein/RNA interactions will be needed to fully understand the translatome and composition of the translational machinery of regenerating ISCs, and the molecular mechanism mediating the role of Aub in mRNA translation in the midgut.

### Aubergine and PIWIL1 in colorectal cancer

Our observations suggest that the upregulation of Aub is a common feature of regenerative and oncogenic ISC hyperproliferation. Alterations of the protein translation machinery is a cancer hallmark.^109^ Over 95% of human CRC cases bear loss of function mutations in the tumour suppressor *APC,* leading to Wnt pathway hyperactivation and the subsequent upregulation of its target gene *myc*.^110^ Myc expression in CRC relies on the activity of the eukaryotic initiation factor eIF4E ^111,112^ and eIF4A1^113^, two factors that, when knocked down, resulted in strong homeostatic stem cell loss in our studies (not shown). Furthermore, eIF3M expression is functionally linked to increased cell proliferation in human colorectal cancer cell lines^114^. It is therefore possible that the mechanisms mediating the upregulation and proliferative function of Aub in intestinal regeneration may extrapolate to the regulation and function of Aub and PIWIL1 in tumours. Our work presents excellent paradigms to address these questions.

### Aubergine functions independently of piRNAs in ISCs of the adult Drosophila midgut

Given that Aub is considered essential for ping-pong piRNAs biogenesis in the germline, our results suggesting its dispensability for the synthesis of the identified ping-pong piRNA-like signature in ISCs were unexpected (**Fig.2**). Detecting a trace amount of contaminated RNA from ovaries, which were not subject to RNAi knockdown in our system, is unlikely because both TE mRNAs and cluster-derived piRNAs are differentially represented in the sorted ISC/EBs compared to ovaries (**Figs.2 and S3**). Instead, this could be explained by either a sufficiency to regulate piRNA with residual levels of Aub expected from RNAi partial gene knockdown or by a non-canonical Piwi and Ago3 dependent regulation of ISC ping-pong piRNAs in the absence of Aub. Alternatively, Aub and Ago3 may work redundantly in ping-pong piRNAs biogenesis in the midgut. Redundant roles of Aub and Ago3 have been reported in the germline.^57^ As such, it would be highly important that work focussing on the regulation and function of piRNAs in ISCs, addresses a potential functional role of Ago3 on its own or redundantly with Aub.

Beyond its role in transposon silencing, Aub has been implicated in several non-canonical processes, including mRNA translation, localisation, and degradation ^115, 102, 116, 117, 69^. In the female germline, Aub regulates the translation of developmental mRNAs to promote stem cell self-renewal and differentiation ^69, 43^. Interestingly, Rojas-Ríos et al. found that the Aub-binding site in *cbl* mRNA closely matches one of the transposon-targeting piRNAs ^43^, whereas Ma et al. reported that Aub iCLIP peaks in more than 1,000 mRNAs showed no sequence homology to piRNAs ^69^. Notably, recent work in mice demonstrated that piRNAs can guide target cleavage even without perfect pairing in the seed region ^118^, likely endowing the germline with a robust genome protection strategy against fast evolving transposable elements. Although similar evidence is lacking in other species, this raises the possibility that Aub may also engage targets through a relaxed, yet still piRNA-dependent, recognition mode.

In contrast to its germline functions, our results show that the ability of Aub to promote intestinal stem cell proliferation and activate Sox21a and Myc expression in the adult midgut appears piRNA-independent as it does not require the piRNA binding or cleavage domain of Aub (**Figs. 3, 4 and S4**). This is reminiscent of PIWIL1 in pancreatic and gastric cancer, which functions independently of its piRNA-binding capacity ^82, 119^. However, while we find that Aub induces protein production in regenerating ISCs, described roles of PIWIL1 role in pancreatic and gastric cancer involve induction of mRNA decay and protein degradation as a co-activator of UPF1 and of the Anaphase-Promoting Complex/Cyclosome respectively ^82, 119^. The unifying principles underlying these non-canonical roles of Aub, and PIWI proteins more broadly, remain unclear. Nevertheless, our findings support the emerging view that PIWI proteins can acquire regulatory functions independent of piRNAs and outside the germline.

### Limitations of the Study

Classical studies characterizing molecular functions of piRNAs and PIWI proteins include immunoprecipitation of PIWI proteins and associated RNAs^6,7,57^.We have not been able to consistently and specifically immunoprecipitate Aub/piRNA complexes from ISCs, likely due to the low abundance of piRNA combined with insufficient amounts of immunoprecipitated Aub from intestinal stem cells when compared to ovary samples. Although the small RNAs detected in our sorted ISCs/EBs preparations showed strong piRNA-like molecular signature, it remains to be determined whether they are indeed associated with PIWI proteins and therefore fit the classical definition of bona-fide piRNAs, and how they may function in ISCs. Such limitations also precluded us from reliably identifying direct target mRNAs or protein partners of Aub in ISCs, as such experiments often require a large amount of material recovered from homogeneous cell populations.^69^ Experimental models amenable for large scale production of cellular material, such as newly emerging intestinal-derived cell lines,^120^ or human intestinal organoids (**Fig.7**), may facilitate conventional biochemical and molecular biology approaches for protein/RNA and protein/protein interaction studies focused on the role of PIWI proteins and piRNAs in the intestine.

## RESOURCE AVAILABILITY

### Lead Contact

- Requests for further information and resources should be directed to and will be fulfilled by the lead contact, Julia B. Cordero.

## Materials Availability

- All unique/stable reagents generated in this study are available from the lead contact without restriction.

## Data and Code Availability

- RNA-seq data have been deposited at GEO at GEO: GSE253621 and GEO: GSE253624, and are publicly available as of the date of publication. TempOSeq RNA seq data from CRC patients has been deposited at NCBI: □PRJNA997336 and are publicly available as of the date of publication. All numeric source data and raw imaging data related to this study has been deposited at Enlighten Research Data Repository at: http://dx.doi.org/10.5525/gla.researchdata.2169 and are publicly available as of the date of publication.
- All original codes has been deposited at Zenodo at https://doi.org/10.5281/zenodo.18608988 and at Github at https://github.com/RippeiHayashi/gut_aubergine and is publicly available as of the date of publication.
- Any additional information required to reanalyze the data reported in this paper is available from the lead contact upon request.

## Acknowledgements

We would like to thank Allison Bardin; Benoit Bîteau; Julius Brennecke; Ginés Morata; Yulii Shidlovskii, Paul Lasko, Marc Amoyel, Sebastian Rampf, Michael Marr, Matthias Hentze and Phillip Zamore for flies and reagents. The Bloomington Drosophila Stock Center; the Vienna Drosophila Resource Center and the *Drosophila* Studies Hybridoma Bank for flies and antibodies. The Core Services and Advanced Technologies at the CRUK Scotland Institute, which is core funded by Cancer Research UK (A31287). With particular thanks to Leo Carlin, Claire Mitchell and Peter Thomason from the Beatson Advanced Imaging Resource, Yi-Hsia Liu and Thomas Gilbey from the FACS facility and Colin Nixon and the Histology Laboratory for assistance with histology. We are grateful to Henri Jasper and Pedro Sousa-Victor for help with cell sorting and mRNA sequencing of midgut ISCs/EBs. We thank all members of the Cordero laboratory for general advice on the project. We thank Donna Markie for SOCCS study coordination and patient recruitment. Human tumour sample analysis was performed on data generated by the TCGA Research Network: https://www.cancer.gov/tcga. The manuscript was critically reviewed by Catherine Winchester (CRUK Scotland 883 Institute). Graphical schematics in the manuscript were created using BioRender under Licence agreement UZ28NVH244.

Work in the Cordero laboratory is funded by Wellcome Trust and Royal Society (104103/Z/14/Z) (223091/Z/21/Z) to J.B.C., core Institutional funds from CRUK to the CRUK Scotland Institute (A31287) and a China Scholarship Council studentship to Y.T. (202006910022). Research in the Hayashi laboratory is supported by the Australian Research Council (DP210102385). The Edwards laboratory is funded by Chief Scientific Office grants (CSO EPD/22/13; K.P. and J.E.) (CSO TCS/22/02; J.E.) and the Scottish Cancer Centre (CTRQQR-2021\100006). Research in the Myant laboratory is supported by the Medical Research Cancer (MRC) MR/X008762/1. Human organoids work was funded by the CRUK Scotland Centre (CTRQQR-2021\100006) and CRUK Program Grant (DRCPGM\100012) to M.G.D. The Gontijo and Heredia laboratory was supported by the FCT (LISBOA-01-0145-FEDER-030753; 10.54499/CEECINST/00102/2018/CP1567/CT0031; 10.54499/DL57/2016/CP1457/CT0016); EXPL/BIA-BID/1524/2021; EXPL/BIA-COM/1296/2021; 10.54499/2022.03859.PTDC, 10.54499/UIDB/04462/2020; 10.54499/UIDB/00329/2020, 10.54499/LA/P/0087/2020; 10.54499/LA/P/0121/2020; LISBOA-01-0145-FEDER-022170; UID/Multi/04462/2019.

## Author contributions

K.B. designed, performed and analysed most of the experiments. L.R.C. initiated the project. Y.T., Y.Y., A.R.C. contributed to *Drosophila* experimental work to the study. K.A.P. and J.E. provided samples, methodology and supervision for TMA analysis. A.B.A., C.V.B., N.D. performed and analysed the work on human intestinal organoids. F.H. and A.M.G. generated *esg-gal4, aub^NH2^* recombinants. F.V.D. and M.G.D. harvested the human tissues from patients and supervised the organoid work. J.P.B. and A.B.A. generated the human organoid lines. K.M. designed, supervised and analysed the work on human intestinal organoids. R.H. designed, performed and analysed piRNA and mRNA sequencing experiments and generated the Aub-GFP transgenic constructs. J.B.C.: study conceptualization, design, supervision, data analysis and funding acquisition. K.B., R.H. and J.B.C. wrote the manuscript with input from other authors.

## STAR METHODS

### KEY RESOURCES TABLE

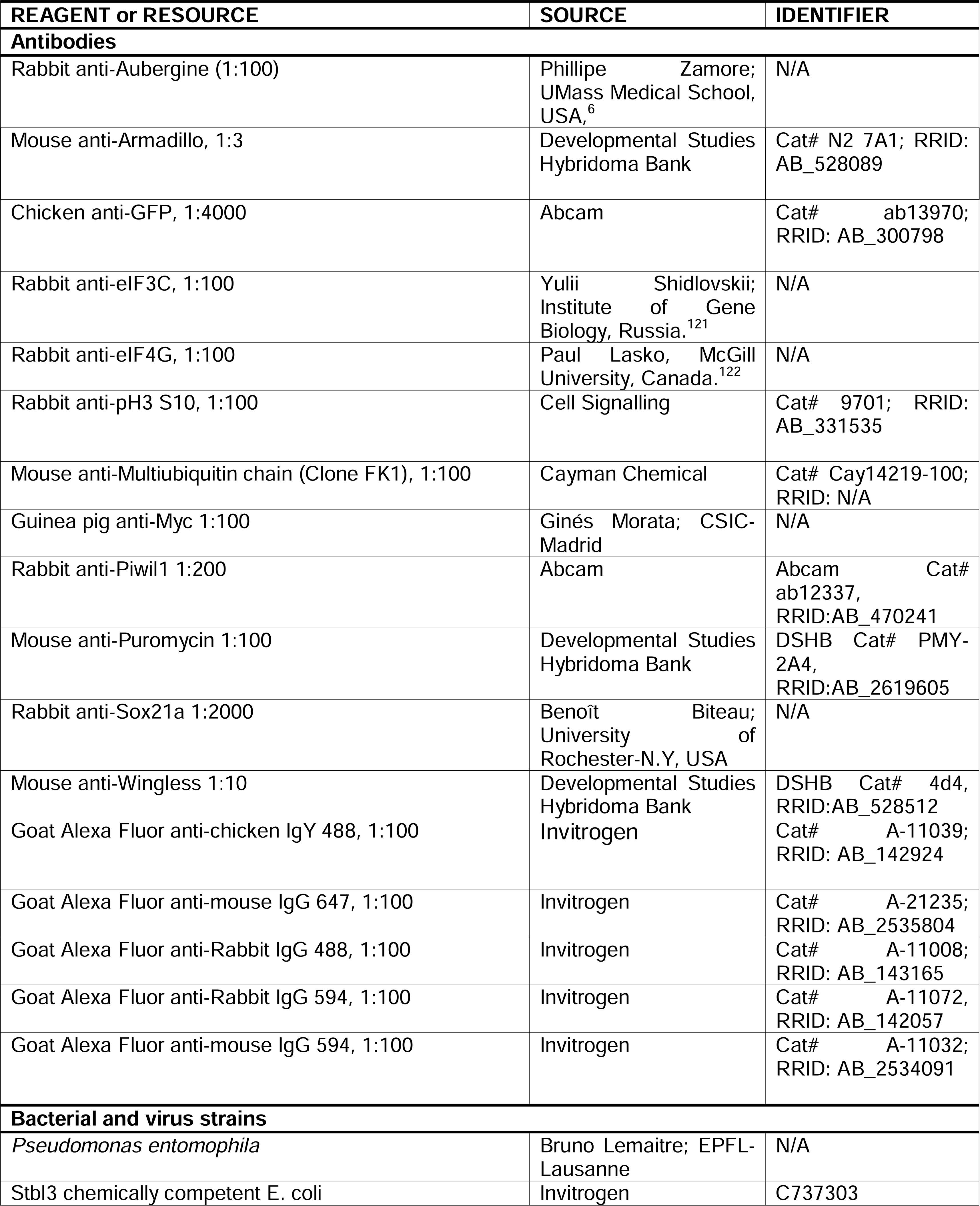
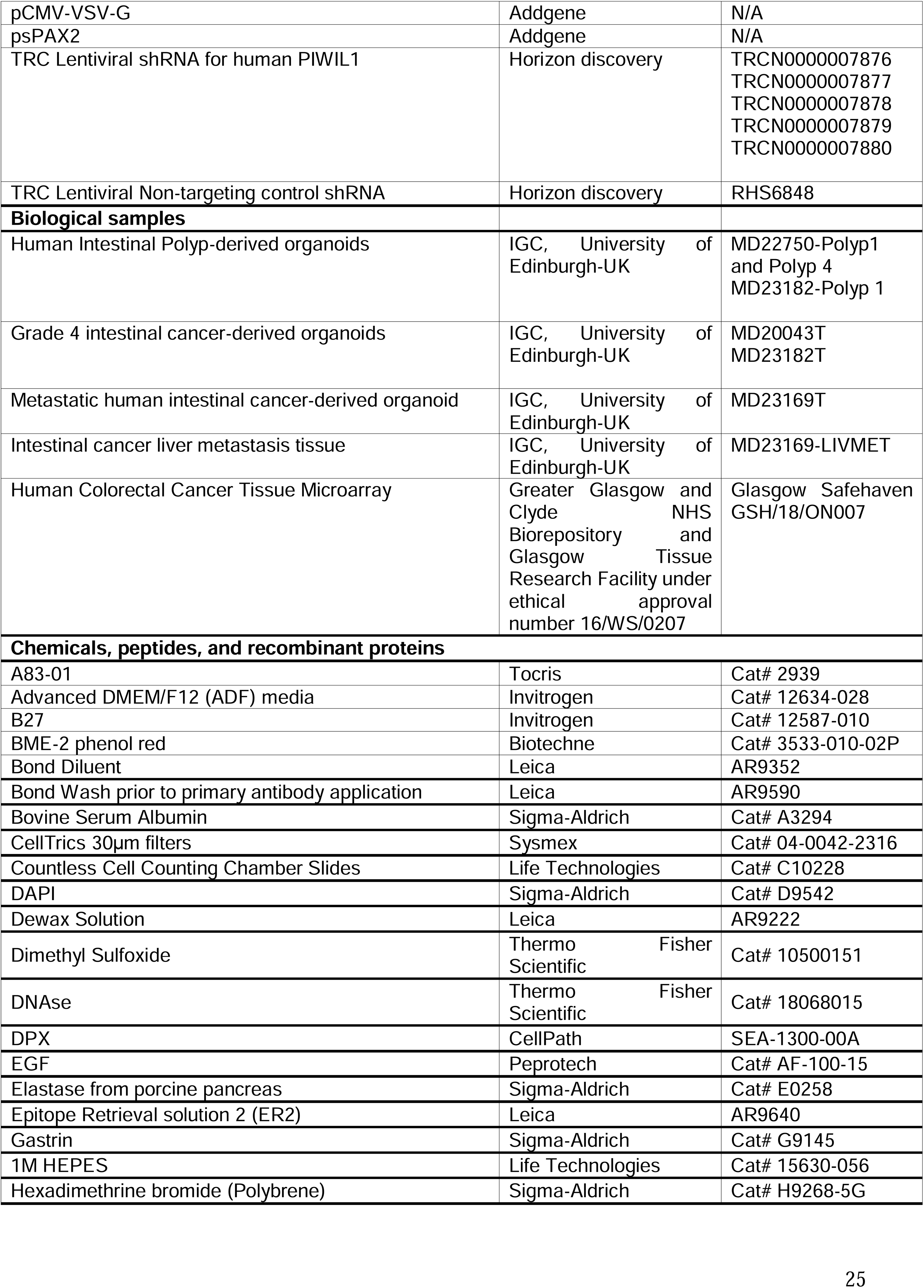
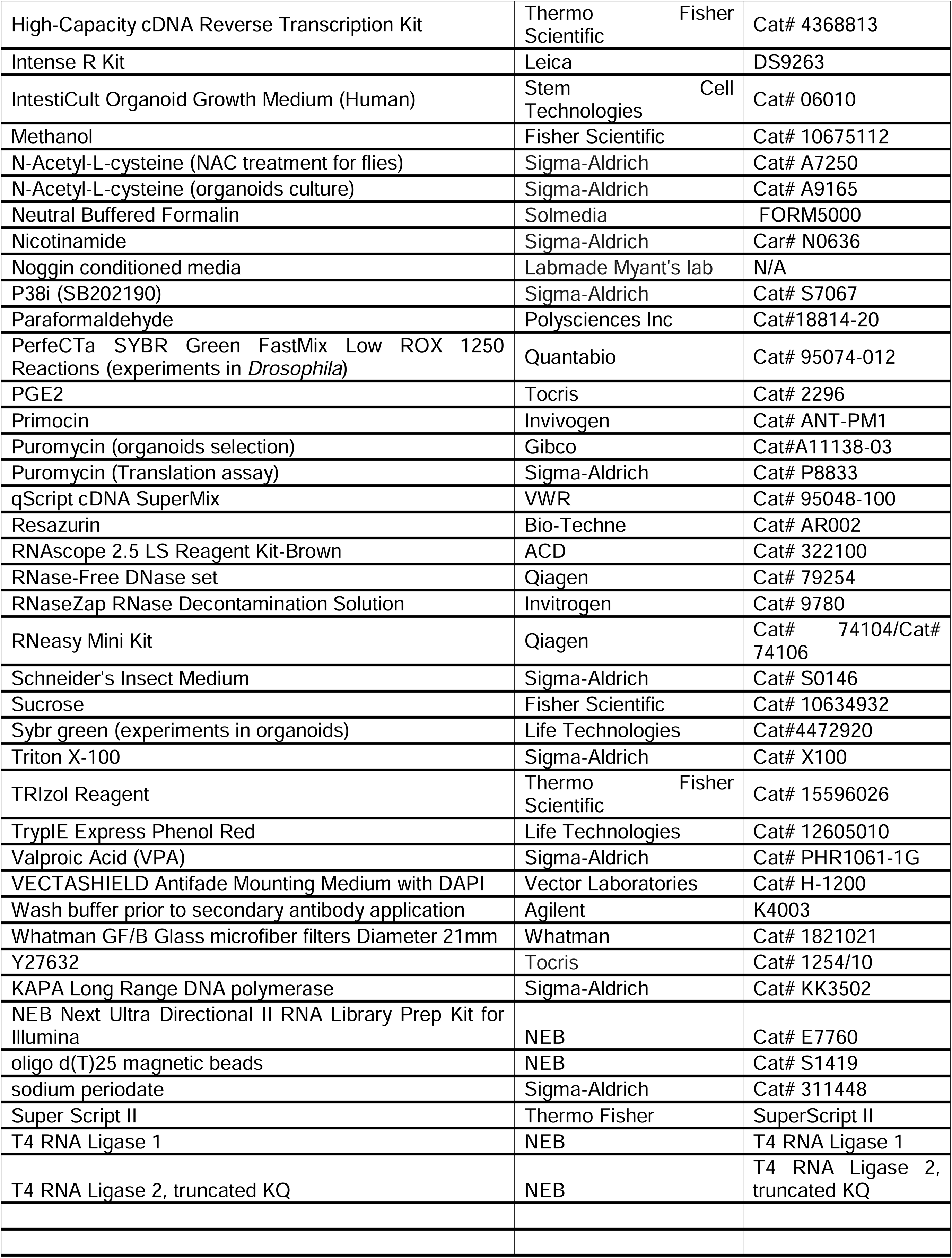
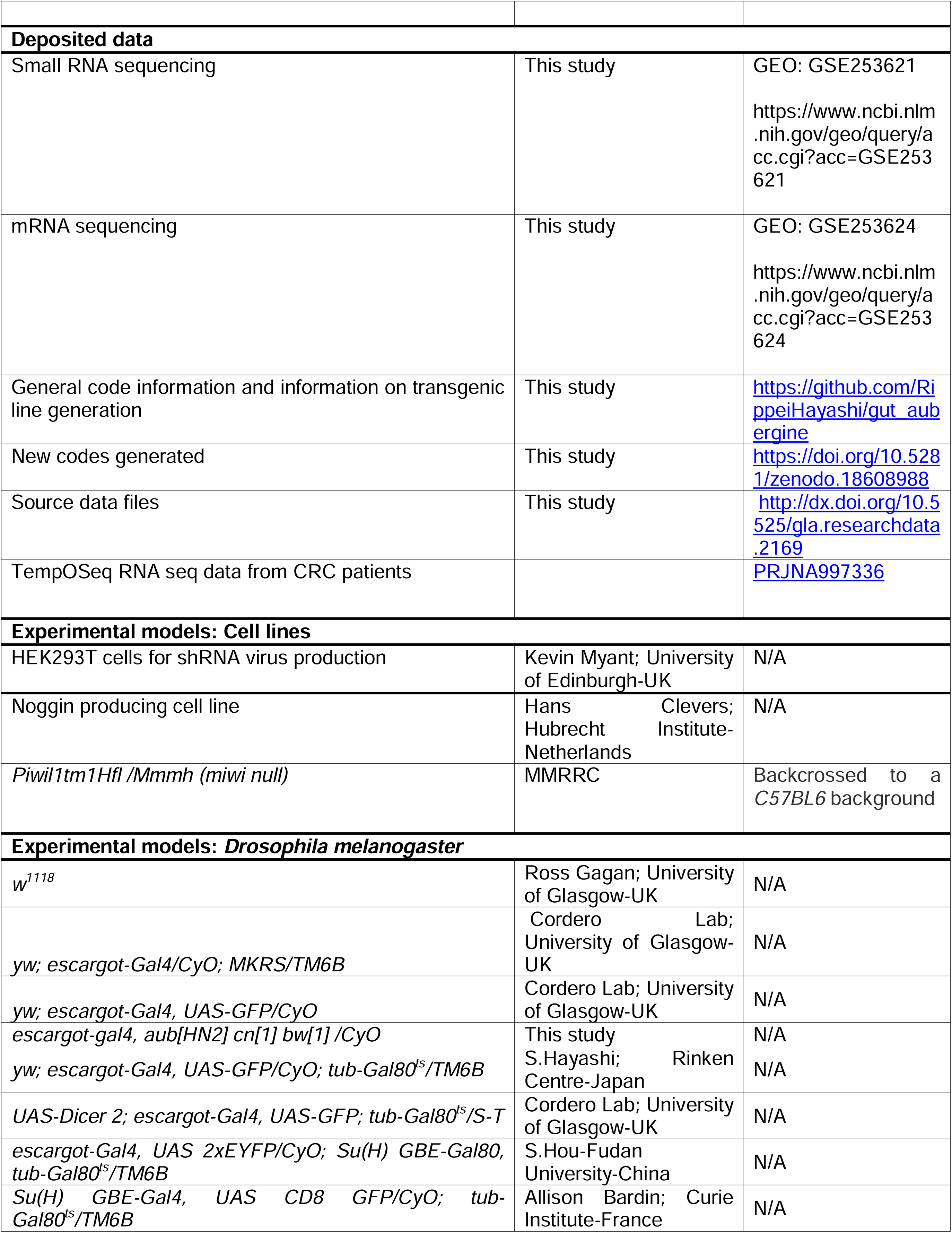
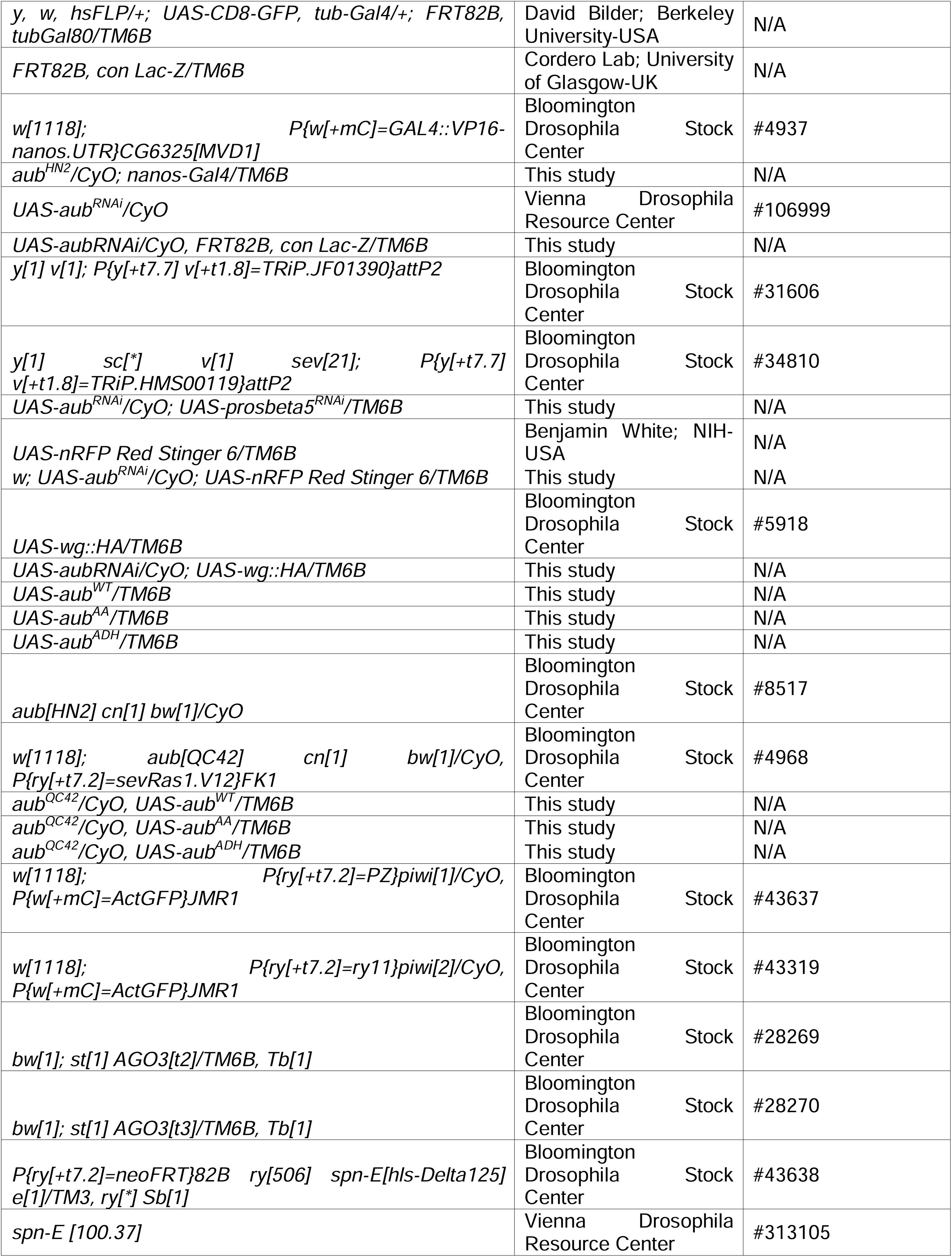
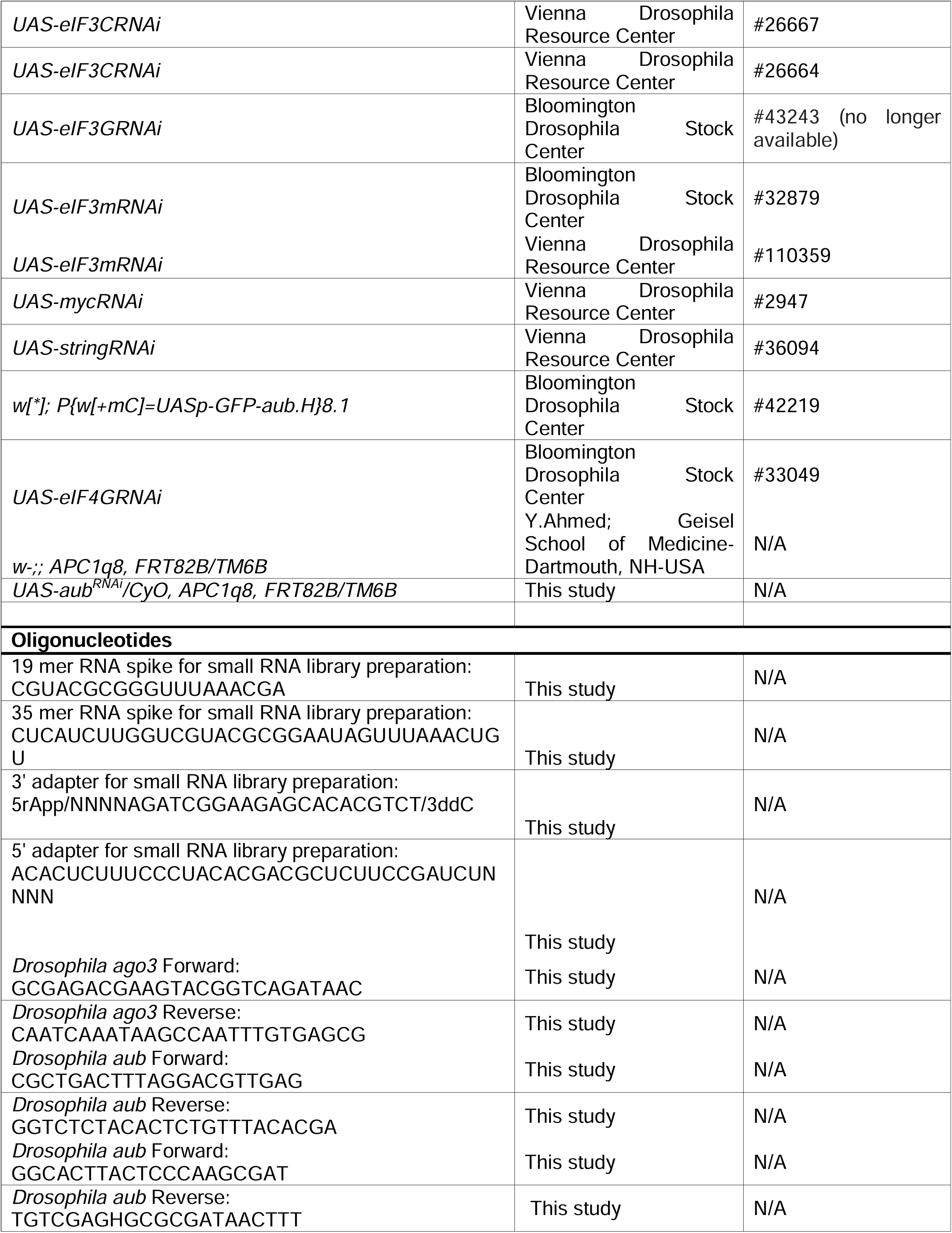
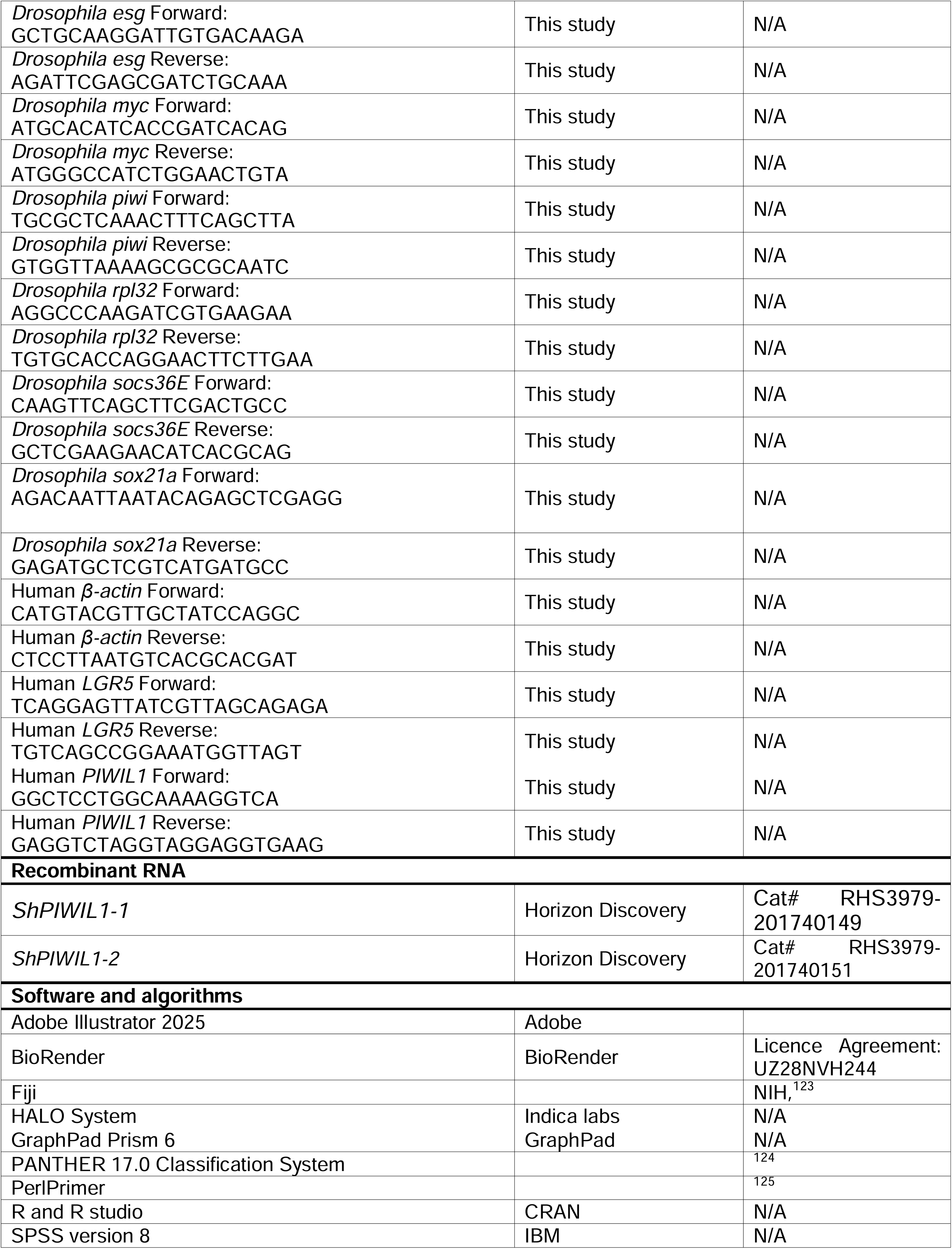
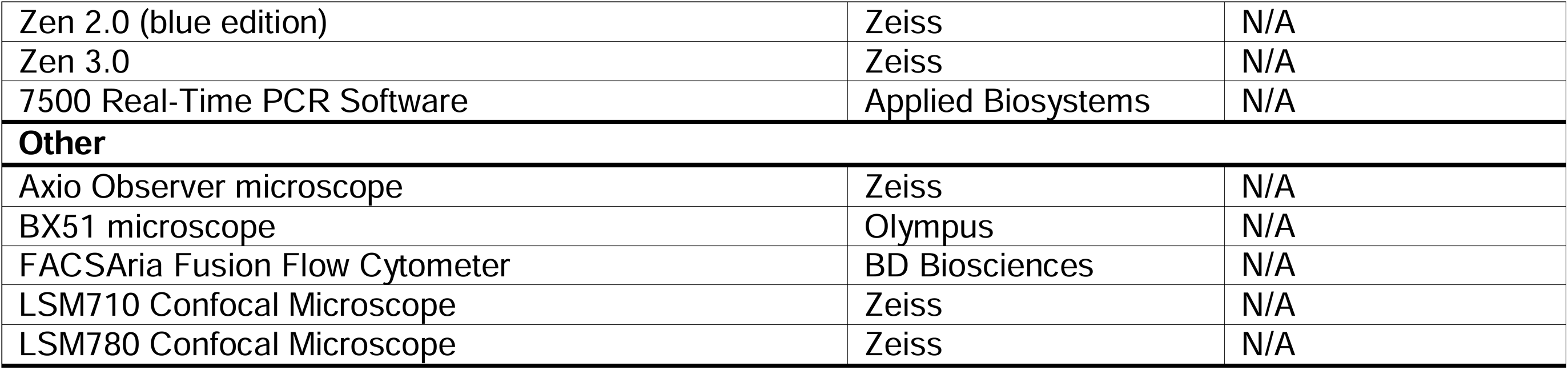

## EXPERIMENTAL MODEL AND STUDY PARTICIPANT DETAILS

### *Drosophila melanogaster* stocks rearing, maintenance and timelines of genetic manipulations

A complete list of fly strains used in this paper is included in the key resources table. Full genotypes for fly lines used in each figure are described in Table S1. Fly stocks were kept in temperature and humidity-controlled incubators set up with a 12h-12h light/dark cycle. For experiments with *aub* mutant lines, crosses were performed at 25°C and the desired F1 progeny was sorted 2/3 days after adult eclosion and kept at 25°C for 5-7 days before dissection. For experiments using the *GAL4*/*GAL80*^ts^ system, flies were crossed at 18°C. The desired F1 progeny was sorted 2/3 days after adult eclosion and was kept at 18°C for 5 days before being transferred to 29°C to allow transgenes activation. In most cases, transgene expression was done for 5-8 days prior to functional experiments. However, in *Aub* gain of function experiments, transgenes were overexpressed for 10 days. Crosses were performed at 25°C and parental lines were flipped twice a week. For MARCM clones, adults of the desired genotype were aged for 3 to 5 days at 25°C before undergoing three 30 minutes heat shocks in one day, in a 37°C water bath. Heat shocked adults were then aged for 7days, 10 days or 14 days at 25°C before dissection. In all cases, experimental flies were flipped every two days on fresh food. Only mated females were used for experiments.

### Human patient information and ethics statement

The human cohort analysed contains 1030 colorectal cancer patients who had undergone elective and potentially curative resection of stage I-III colorectal adenocarcinoma in a single surgical unit at the Glasgow Royal Infirmary, Western Infirmary or Stobhill Hospitals (Glasgow, UK) between 1997 and 2007. Resection was considered curative based on pre-operative computed tomography and intra-operative findings. The data are stored within the Glasgow Safehaven (GSH21ON009) and ethical approval was in place for the study (MREC/01/0/36).

### Human patient-derived organoids (PDOs)

The human derived PDOs used in this study were generated in the Edinburgh component of the CRUK Scotland Centre under the supervision and direction of MGD and FVND, University of Edinburgh: MD20043 is an 81-year-old male with Stage 4 rectal cancer, (T3N2M1). MD23169 is male with Stage 4 ascending colon tumour along with current liver metastasis (T3N1M1). Ethical approval for human CRC organoid derivation was carried out under NHS Lothian Ethical Approval Scottish Colorectal Cancer Genetic Susceptibility Study 3 (SOCCS3) (REC reference: 11/SS/0109). All patients provided fully informed consent for analysis of tumour and normal tissue that was surplus to requirements for pathological assessment, clinical annotation and generation of organoids.

## METHOD DETAILS

### Human organoid culture media and shPIWIL1 knockdown

Human carcinoma organoids were cultured in PDO media containing 1% Noggin conditioned media, 1x B27, 50□ng/mL EGF, 10□nM Gastrin, 10□nM PGE2, 10□mM Nicotinamide, 10□µM SB202190, 600□nM A83-01, 12.5 mM N-Acetylcysteine in ADF (with 1X PenStrep, HEPES, Glutamine, 100µg/mL Primocin).

For transduction conditions, human organoids were pre-treated with IntestiCult organoid growth media (OGM) and VPA (1:10,000) for 48h after 2 days post-split (Day-1). On transduction day (Day1), human organoids were digested into single-cell suspension in TrypLE with 10□µM Y27632 for 8□min at 37□°C with mechanical dissociation every 4□min. Single cell suspension (with 8ug/ml Polybrene) were combined with viral particles containing, TRC non-targeting (NT) control, TRC shPIWIL1-1 (76), and shPIWIL1-2 (78) and placed on BME layer. On Day2, media was changed into IntestiCult OGM + VPA (1:10,000) + 10□µM Y27632. On Day3, antibiotic selection started with 2µg/mL Puromycin in IntestiCult OGM. On Day6, RNA and protein samples are collected, and colony formation ability (clonogenicity) assay started. For clonogenicity, organoids from Day 6 were collected, washed once or twice with ice cold PBS. Then, human organoids were digested into single-cell suspension in TrypLE with 10□µM Y27632 for 5-8□min at 37□°C with mechanical dissociation every 3-4□min. 2000 single cells in 5µl BME drops per well were plated and IntestiCult OGM with 2µg/mL Puromycin added after 15 min to set BME. Organoid formation was observed throughout the clonogenicity assay. On Day 14, organoid numbers were counted manually on Leica Light microscope and pictures were taken on EVOS FL Cell Imaging System. For cell viability, the media was replaced with fresh media containing 10% Resazurin for 24h in 37°C incubator. The next day, relative cell viability was measured with a TECAN Spark microplate reader in 96-well plate with three technical replicates. IntestiCult OGM media was used exclusively during transduction and the single-cell stage of organoid development for clonogenicity assays. Otherwise, only PDO media was utilized for maintaining the organoids.

### Damage-induced intestinal regeneration

To induce intestinal regeneration, mated female flies were fed on filters (Whatman) soaked in 5% sucrose only (control condition) or with the pathogenic bacteria *Pseudomonas entomophila* (*Pe*) at OD_600_=25 (damage condition) for 16 hours prior to dissection. The bacterial solution was obtained after overnight culture in LB medium at 29°C under 220 rpm shaking overnight, after which bacteria were pelleted (Beckman Coulter JS-4.2 rotor, 10 min, 4000 rpm) and diluted in a solution of 5% sucrose to reach OD_600_=25.

### Immunohistochemistry

#### Drosophila tissue

The dissection of guts was performed in PBS at room temperature and 16 hours after feeding the mated female flies with sucrose or *Pe*. After dissection, guts were immediately fixed in solution of 4% paraformaldehyde (Polysciences Inc) for 1 hour. Tissues were then washed in PBST (PBS with Triton 0.2%) before incubation overnight at 4°C with the primary antibodies diluted in blocking solution PBT (with 0.5% BSA). The day after, guts were washed 3x15 minutes with PBST at room temperature before incubation with the secondary antibodies for 2 hours at room temperature. Guts were then washed 3x15 minutes with PBST at room temperature before mounting in DAPI containing mounting media. Slides were kept at 4°C before imaging. To improve Arm and Aub staining, guts were placed in methanol for 10 minutes at room temperature after the fixation step with 4% paraformaldehyde.

#### Human and mouse tissue

Immunohistochemistry (IHC) staining was performed on a previously constructed human tissue microarray or on 4µm formalin fixed paraffin embedded sections, which had previously been incubated in a 60□C oven for 2 hours. Tissue slides for PIWIL1 detection were stained on a Leica Bond Rx Automated Stainer. Samples underwent on-board dewaxing and epitope retrieval using ER2 solution (Leica) for 20 minutes at 95°C. Sections were rinsed with Leica wash buffer before peroxidase block was performed using an Intense R kit (Leica) for 5 minutes. Sections were rinsed with wash buffer (Leica) before anti-PIWIL1 application at an optimal dilution (1/200). The sections were then rinsed with wash buffer (Agilent) for 30 minutes, before secondary antibody application. The sections were rinsed with wash buffer, visualized using DAB and counterstained with Haematoxylin in the Intense R kit. To complete the IHC staining sections were rinsed in tap water, dehydrated through a series of graded alcohols, and placed in xylene. The stained sections were cover slipped in xylene using DPX mounting.

### NAC treatment

Flies were placed in empty vials with filters soaked with a 5% sucrose solution alone or containing 20□mM NAC 48h prior dissection. Flies were then fed with either sucrose; *Pe* (OD_600_=25); 5% sucrose□+□20□mM NAC or 5% sucrose□+□20□mM NAC□+□*Pe* (OD_600_=25) 16h prior dissection.

### Generation of *aub* transgenes

The three pUASt-attB-GFP constructs of *aub* were generated for this study. A complementary DNA (cDNA) of *aub* was amplified from *w^1118^* flies and cloned into the pUASt-attB plasmid, that was originally made by the Basler group (GenBank accession EF362409)^126^ and received as a gift from Julius Brennecke. A GFP cDNA was inserted N-terminally to *aub* cDNA. Standard site-directed mutagenesis was used to introduce amino acid substitutions to generate *aub^AA^* (Y345A/Y346A,^59^) and *aub^ADH^* (D636A,^19^). Plasmid construction and sequences are available at https://github.com/RippeiHayashi/gut_aubergine.

### Egg laying/eclosion assay

Crosses to obtain experimental animals were kept at 18□C. Virgin females of the desired genotype were selected and placed into fresh vials and allowed to mate with *w^1118^* males. A maximum of 5 females and 5 males were housed in each vial and allowed to mate over a period of 24 hours at 18□C. The following day, males were removed, and females were moved onto fresh food and incubated at 25□C. Females were flipped into new vials every 24 hours. The number of eggs in each vial was counted and recorded at the same time. This process was repeated 3 more times. Importantly, the number of live females was recorded each time, at the time of flipping. Any females that died during the night were included in the eggs/adult ratio but were excluded from the subsequent recordings. Three replicates of these experiments were performed.

### Protein translation assay

For general protein translation assessment, flies were fed on filters (Whatman) soaked in 5% sucrose only or with *Pe* OD_600_=25 16 hours prior to gut dissection in Schneider’s insect medium (Sigma-Aldrich) containing Puromycin (stock solution 25 mg/mL, Sigma-Aldrich) with a final concentration of 25µg (1μL of stock solution) of puromycin in 5mL of Schneider’s insect medium. During dissection, Malpighian tubules and crop were kept attached to the gut and precaution was taken to keep the intestines intact. Guts were then incubated at room temperature in puromycin containing medium for 45 minutes. After incubation, standard immunochemistry protocols were applied.

### *Drosophila* protein expression time course

Flies were fed with fed on filters soaked in 5% sucrose only *Pe* (OD_600_=25). Half of the flies were dissected following 16h post-infection. The second half was flipped on regular food and dissected 24h later (recovery).

### Fluorescence-activated single cell sorting

Control flies and flies expressing RNAi against *aub* were collected between 2 to 3 days after hatching and kept at 18°C for 5 days before being transferred to 29°C. Flies were kept are 29°C for 7 days and were flipped into fresh food vials every two days. Fly guts were dissected in cold PBS 16 hours after animals being fed with 5% sucrose only or 5% sucrose containing *Pe*. The crop, the Malpighian tubules and the hindgut were carefully removed. Midguts were then collected and transferred in an Eppendorf containing 400 µL of PBS and kept on ice. Each Eppendorf contained roughly 100 midguts. Once all the tissue was dissected, 10 µL of elastase (Sigma-Aldrich, 10 µg/ µL) was added to each Eppendorf. Samples were kept at 27°C in a heat block for at least 1 hour until midguts were fully dissociated. Tissues were further disrupted by pipetting up and down every 5 minutes. Samples were then centrifuged at 300 rcf for 20 minutes at 4°C. After removing the supernatant, the pellet was resuspended in cold PBS. To eliminate enterocytes as much as possible, the cell suspension was filtered with a 30 µm filter (Sysmex). 500 µL of cold PBS were added to wash the filter. After filtration, samples were ready for sorting. Sorting was performed by the CRUK Scotland Institute flow cytometry facility on a FACSAria Fusion Flow Cytometer (BD Biosciences). GFP or Red stinger positive controls were used to set up the flow cytometer parameters that were used for all the conditions. In parallel to the sorting, a few microliters of samples were taken and stained with 20 µL of DAPI to visualize cell death. Between 250 and 500 flies were dissected per condition, resulting in 250,000 to 750,000 sorted cells. Three biological replicates were dissected for each condition. Sorted cells were pelleted by centrifugation at 300 rcf for 20 minutes at 4°C. After carefully removing the supernatant, 800 μL of TRIzol (Invitrogen) was added to the samples, followed by RNA extraction and sequencing.

### Small RNA sequencing

Small RNA sequencing was performed on either whole midguts, ovaries or sorted ISCs/EBs. For small RNA preparation from whole midguts, flies were dissected in cold PBS 16 hours after being fed with 5% sucrose only or 5% sucrose containing *Pe*. The crop, the Malpighian tubules and the hindgut were carefully removed. Three replicates of 40 midguts each were dissected for each condition. Guts were then collected and transferred to an Eppendorf containing 400 µL of PBS and kept on ice. Once all the tissue was dissected, 10 µL of elastase (Sigma-Aldrich, 10 µg/µL) was added to each Eppendorf. Samples were kept at 27°C in a heat block for at least 1 hour until the guts were totally degraded. Tissues were disrupted by pipetting up and down every 5 minutes. Samples were then centrifuged at 300 rcf for 20 minutes at 4°C. After removing the supernatant, the pellet was resuspended in 800 µL of TRIzol (Invitrogen). The same protocol was applied for the ovaries. Approximately 10 pairs of ovaries were dissected. We generated small RNA libraries from 1 ∼ 5 μg of total RNA using a modified protocol from the original method.^127^ 19 to 35 nucleotides-long RNA was first selected by PAGE using radio-labelled 19mer spike (5′-CGUACGCGGGUUUAAACGA) and 35mer spike (5′-CUCAUCUUGGUCGUACGCGGAAUAGUUUAAACUGU). The size-selected RNA was precipitated, oxidised by sodium periodate,^128^ and size-selected for the second time by PAGE. The size-selected oxidised small RNAs were ligated to the 3’ adapter from IDT (5rApp/NNNNAGATCGGAAGAGCACACGTCT/3ddC where Ns are randomised) using the truncated T4 RNA Ligase 2, truncated KQ (NEB), followed by a third PAGE to remove non-ligated adapters. Subsequently, the RNA was ligated to the 5’ adapter from IDT (ACACUCUUUCCCUACACGACGCUCUUCCGAUCUNNNN where Ns are randomised) using the T4 RNA Ligase 1 (NEB). Adapter-ligated RNA was reverse-transcribed by SuperScript II (Thermo Fisher) and amplified by KAPA LongRange DNA polymerase (Sigma, KK3502) using the universal forward primer, Solexa_PCR-fw: (5′- AATGATACGGCGACCACCGAGATCTACACTCTTTCCCTACACGACGCTCTTCCG ATCT) and the barcode-containing reverse primer TruSeq_IDX: (5′-CAAGCAGAAGACGGCATACGAGATxxxxxxGTGACTGGAGTTCAGACGTGTGCTC TTCCGATCT where xxxxxx is the reverse-complemented barcode sequence). Amplified libraries were multiplexed and sequenced on a HiSeq platform in the paired-end 150 bp mode by GENEWIZ/Azenta.

### Small RNA sequencing analysis

The R1 sequencing reads were trimmed of the Illumina-adapter sequence using the FASTX-Toolkit from the Hannon Lab (CRUK Cambridge Institute). The 4 random nucleotides at either ends of the read were further removed. The trimmed reads of 18 to 40 nt in size were first mapped to the infrastructural RNAs, including ribosomal RNAs, small nucleolar RNAs (snRNAs), small nuclear RNAs (snoRNAs), microRNAs, and transfer RNAs (tRNAs) using Bowtie 1.2.3 allowing up to one mismatch. Sequences annotated in the dm6 r6.31 assembly of the *Drosophila melanogaster* genome were used. The trimmed and unfiltered reads were then mapped to the dm6 genome using Bowtie allowing up to one mismatch. Reads that uniquely mapped to the genome were analysed for the tile analysis. Reads that mapped to the 100 nt upstream and downstream genomic regions of tRNA, snRNA and snoRNA insertions as well as those that mapped to the *aubergine* gene locus were removed from the tile analysis. Bedtools 2.28.0 was used to count the coverage of the mapped reads. Endogenous siRNA reads (annotated as hpRNA) were used for normalisation. The trimmed and unfiltered reads were separately mapped to the curated sequences of *Drosophila melanogaster* transposons,^57^ using Bowtie allowing up to three mismatches with the option of --all --best --strata. The size distribution of the transposon-mapping reads was made using ggplot2 3.4.4 in R 4.0.0. Nucleotide frequencies around the first nucleotide position of antisense and the tenth nucleotide position of the sense transposon-mapping piRNA reads (longer than 22 nucleotides) were counted and visualised using weblogo 3.7.8. Frequencies were measured in the window of 11 nucleotides, and the z scores were calculated as the deviation of the frequency value at the first (antisense reads) and tenth (sense reads) positions from the mean frequency divided by the standard deviation of the frequencies. Small RNA sequencing libraries from Siudeja et al.^56^ were analysed in the same way except for using the adapter sequence TGGAATTCTCGGGTGCCAAG and not trimming the 4 nucleotides at either end.

### PolyA-selected RNA sequencing

Polyadenylated RNA was purified from the DNase-treated total RNA using the oligo d(T)25 magnetic beads (NEB, S1419) and used for the library preparation. Libraries were cloned using the NEBNext Ultra Directional II RNA Library Prep Kit for Illumina (NEB, E7760), following the manufacturer’s instruction, and amplified by KAPA polymerase using the same primers as for the small RNA sequencing.

### PolyA-selected RNA analysis

Both R1 and R2 reads from the polyA-selected RNA sequencing reads were trimmed of the Illumina-adapter sequences using the FASTX-Toolkit. The trimmed reads were subsequently filtered by the sequencing quality. Only the paired and unfiltered reads were then mapped to the dm6 r6.31 transcriptome combined with curated sequences of *Drosophila melanogaster* transposons using salmon/1.1.0 with the options of --validateMappings --incompatPrior 0.0 --seqBias --gcBias. Length-normalised transposon mRNA reads per one million transcripts including host mRNAs were compared between different libraries. polyA-selected RNA sequencing libraries from ovarian samples from Senti et al.^57^ were also analysed in the same way.

### RNA extraction and RT-qPCR

#### Drosophila tissues

Total RNA was extracted from tissues dissected in cold PBS. Three or four replicates of 20 to 40 guts were dissected for each condition. Depending on the experiments, sorted ISCs/EBs cells, R4-R5 regions, ovaries and/or whole guts were homogenised in 100 µl of Trizol (Thermo Fischer Scientific) using a mortar and a pestle first, followed by adding further 700 µl of Trizol. 200 µl of Chloroform was added and mixed well with Trizol by vortexing before centrifugation on a bench top centrifuge. The aqueous phase was saved, and RNA was precipitated with iso-propanol. RNA was treated with DNase (Thermo Fischer Scientific) and quantified using a NanoDrop Spectrophotometer. About 1 µg of RNA was converted to cDNA using a high-capacity cDNA reverse transcription kit (Thermo Fisher Scientific). RT-qPCR was performed using Perfecta SYBR green fast mix (Quantabio) as per the manufacturer’s instructions on an Applied Biosystems QuantStudio 3 fast real-time PCR system and each reaction was carried out in triplicates. A standard curve was created using 1:10 dilutions from pooled cDNA samples. The amount of sample was extrapolated from this standard curve and normalised using data from the housekeeping gene *rpl32*. Melting curves were conducted priorly to ensure only one product was formed from each pair of primers. Data was exported and analysed in excel. The results are shown as the ratio of average of mRNA levels of the candidate gene/*rpl32*.

#### Intestinal organoids

RNA samples from 3 biological replicates were isolated by using RNeasy Mini Kit (Qiagen #74106) following manufacturer’s protocol. RNA samples were subsequently treated with RNase-Free DNase Set (Qiagen, #79254). RNA samples were analysed for quality and quantity using Thermo Scientific™ Nanodrop™ (Thermo Fisher). 500ng RNA used for cDNA generation using qScript™ cDNA SuperMix (VWR, #95048-100). The results are shown as the ratio of average mRNA levels of the candidate gene/β*-actin*.

### TCGA analysis

To study *PIWIL1* RNA expression across the different CMS subtypes of CRC or according to the microsatellite status of the patients, we took advantage of the TCGA datasets from the PanCancer Atlas. Analysis has been performed on open access data from colonic adenocarcinoma patients only. Differential gene expression was then assessed, and statistics have been performed using Graphpad.

### Samples visualization and image acquisition

PH3 counting was performed using a BX51 Olympus. Fluorescence images of *Drosophila* intestines were acquired using a confocal Zeiss 710 or a Zeiss LSM 780. Stained TMAs slides were scanned and analysed using the HALO system (indica labs).

## QUANTIFICATION AND STATISTICAL ANALYSIS

### Immunofluorescence staining

Quantification of ISC proliferation was done by manually counting the number of PH3-positive cells per posterior midguts. Quantification of Arm, eIF3C, eIF4G, Multiubiquitination, Myc, Puromycin, Sox21a and Wg staining was performed by measuring the mean fluorescence intensity of the protein in all GFP positive cells on a SUM or MAX projection generated via Fiji. Prior the measurement, a mask was created via Fiji to delineate the GFP positive cells only, using the tool “Create a selection”. In parallel, mean fluorescence intensity of the background was measured on a specific region of interest on each SUM projection and subtracted from the signal measured in the cells.

Quantification of Aub staining was performed by measuring the mean fluorescence intensity per cell on a SUM projection done by Fiji. This measure was blindly performed in three different GFP positive cells per gut for Aub. In parallel, mean fluorescence intensity of the background was measured on a specific region of interest on each SUM projection and subtracted from the signal measured in the cell. This quantification was repeated in 5 different guts per biological replicate.

In the rescue experiments (*aub^WT^, aub^AA^* and *aub^ADH^*), GFP and Aub staining were quantified by measuring the mean fluorescence intensity of the protein in all GFP positive cells on a MAX projection realised via Fiji. Prior the measurement, a mask was created on Fiji to delineate the GFP positive cells only via using the tool “Create a selection”. In parallel, mean fluorescence intensity of the background was measured on a specific region of interest on each SUM projection and subtracted from the signal measured in the cells. Quantification of the % of GFP positive area was measured using the same mask and divided by the gut area, measured with the DAPI staining.

### MARCM clones

Quantification of ISC proliferation in MARCM clones was performed by manually counting the number of DAPI positive cells in each GFP-positive clone. The statistics were done with GraphPad Prism 6. The tests used for each experiment are described in the figure legends.

### Transcriptional profiling of patient tissue

The Glasgow cohort consisted of n=787 stage 1-3 CRC patients who underwent surgery with curative intent within Greater Glasgow and Clyde between 1997 and 2013. Formalin fixed paraffin embedded tumour resections were annotated for epithelial rich regions. These areas were extracted and profiled for full transcriptome expression using TemOSeq as previously described.^129^ Patients were excluded from analysis due to mortality within 30 days of surgery. Raw gene counts were normalised using DESeq2 in RStudio. *PIWIL1* expression was assessed for association with TNM stage via Kruskal Wallis testing in GraphPad prism version 10. Data were dichotomised into high and low expression groups using the *Survminer* package in R Studio. Kaplan Meier survival analysis was performed using the *Survival* package in R Studio. This study was approved by the Research Ethics Committee of the West Glasgow University Hospitals NHS Trust (NHS GG&C REC ref. 22/WS/0020), in accordance with Human Tissue (Scotland) Act 2006, which included policy on consent. Data were deposited and accessible within Glasgow Safehaven (GSH21ON009).

### Statistics

Detailed statistical information for each experiment within the study is provided within the figure legends. In all cases data represent the mean□±□SD. ns, not significant (*P*□>□0.05); \**P*□<□0.05, \*\**P*□*<*□0.01, \*\*\**P*□<□0.001 and \*\*\*\**P*□*<*□0.0001.

## Supplemental information

Supplementary Data: Figures S1-S7.

Table S1: Full genotypes used in this study. Related to STAR Methods.

