## Supplementary data file for "Non-gonadal PIWI protein, Aubergine, regulates regenerative stem cell proliferation and tumorigenesis in the *Drosophila* adult intestine"

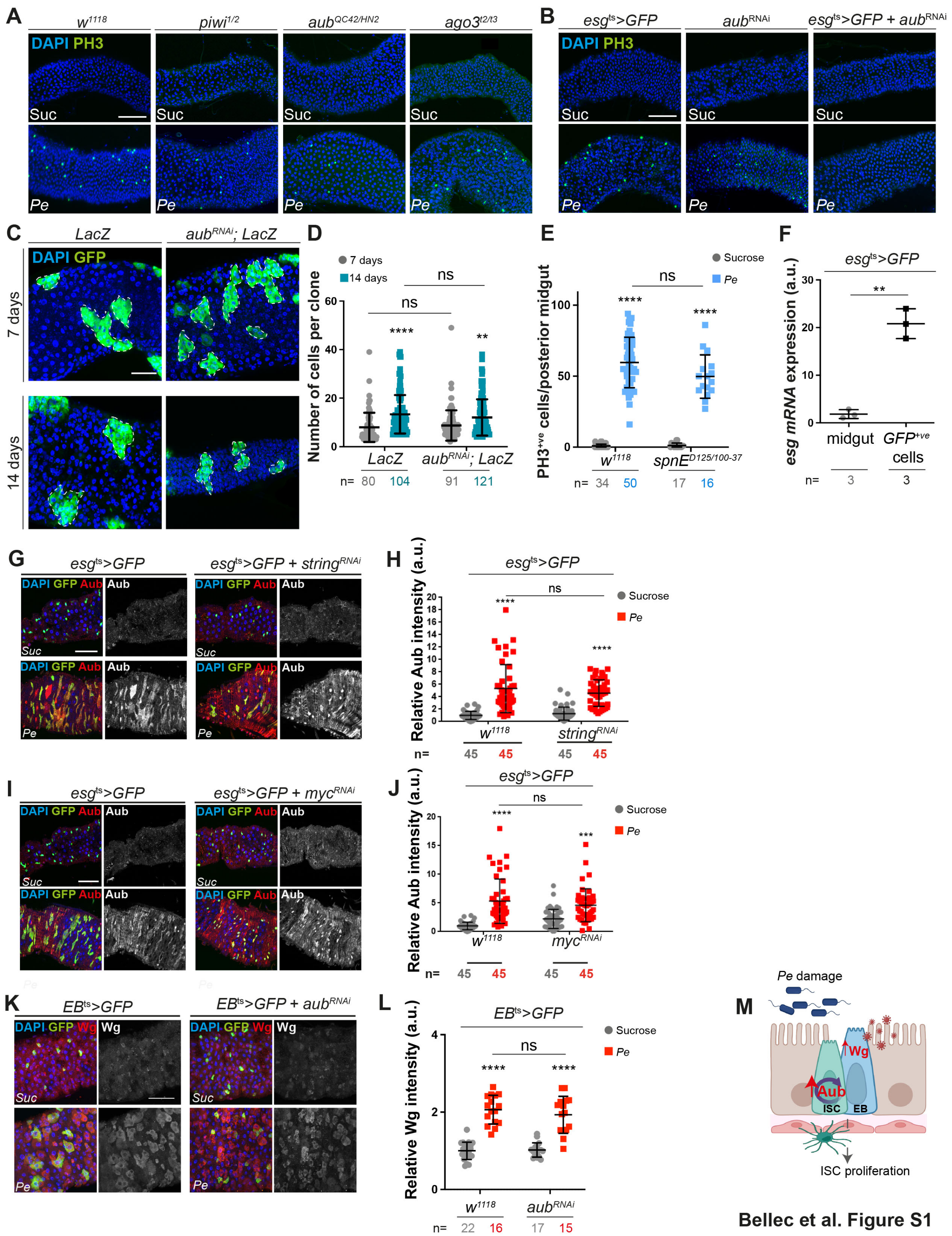

**Figure S1: Aub regulates adult intestinal regeneration in *Drosophila.* Related to Figure 1. (A)** PH3 staining (green) in the posterior midguts from $w^{1118}$ flies or from whole mutant flies fed with sucrose or *Pe*. Scale bar = 100µm. **(B)** PH3 staining (green) in midguts of flies expressing GFP, $aub^{RNAi}$ alone or $aub^{RNAi}$ within ISCs/EBs and fed with sucrose or *Pe*. Scale bar =100µm. **(C)** MARCM clones (green) expressing a *LacZ* transgene only or with $aub^{RNAi}$. Dashed white lines delineate clonal margins. **(D)** Quantification of clone size as in C. n = number of clones. **(E)** PH3-positive cells in the posterior midguts **(F)** Relative mRNA expression of *esg* in whole guts or in sorted ISCs/EBs. n=biological replicates. T test. **(G)** Aub staining (red and grey) in the posterior midguts of flies expressing $esg^{ts}$*>GFP* or $esg^{ts}$*>GFP* + $string^{RNAi}$ for cell cycle inhibition. **(H)** Quantification of staining as in G. **(I)** Aub staining (red and grey) in $esg^{ts}$*>GFP* or $esg^{ts}$*>GFP* + $myc^{RNAi}$ midguts to block ISC proliferation. **(J)** Quantification of staining as in I. **(K)** Wg staining (red) in the posterior midguts of flies expressing GFP (green) only or with $aub^{RNAi}$ in EBs and fed with sucrose or *Pe*. **(L)** Quantification of staining in as in K. Nuclei are identified with DAPI. Scale bars = 50µm. **(M)** Scheme of Aub and Wg regulation within the intestinal epithelium of *Drosophila*. Unless otherwise noted, two-way ANOVA followed by Sidak's multiple comparisons tests were used for statistical analysis, and n = number of midguts/flies quantified. a.u., arbitrary units. Data are represented as mean +/- SD. ns, not significant; *$P < 0.05$, **$P < 0.01$, ***$P < 0.001$; ****$P < 0.0001$.

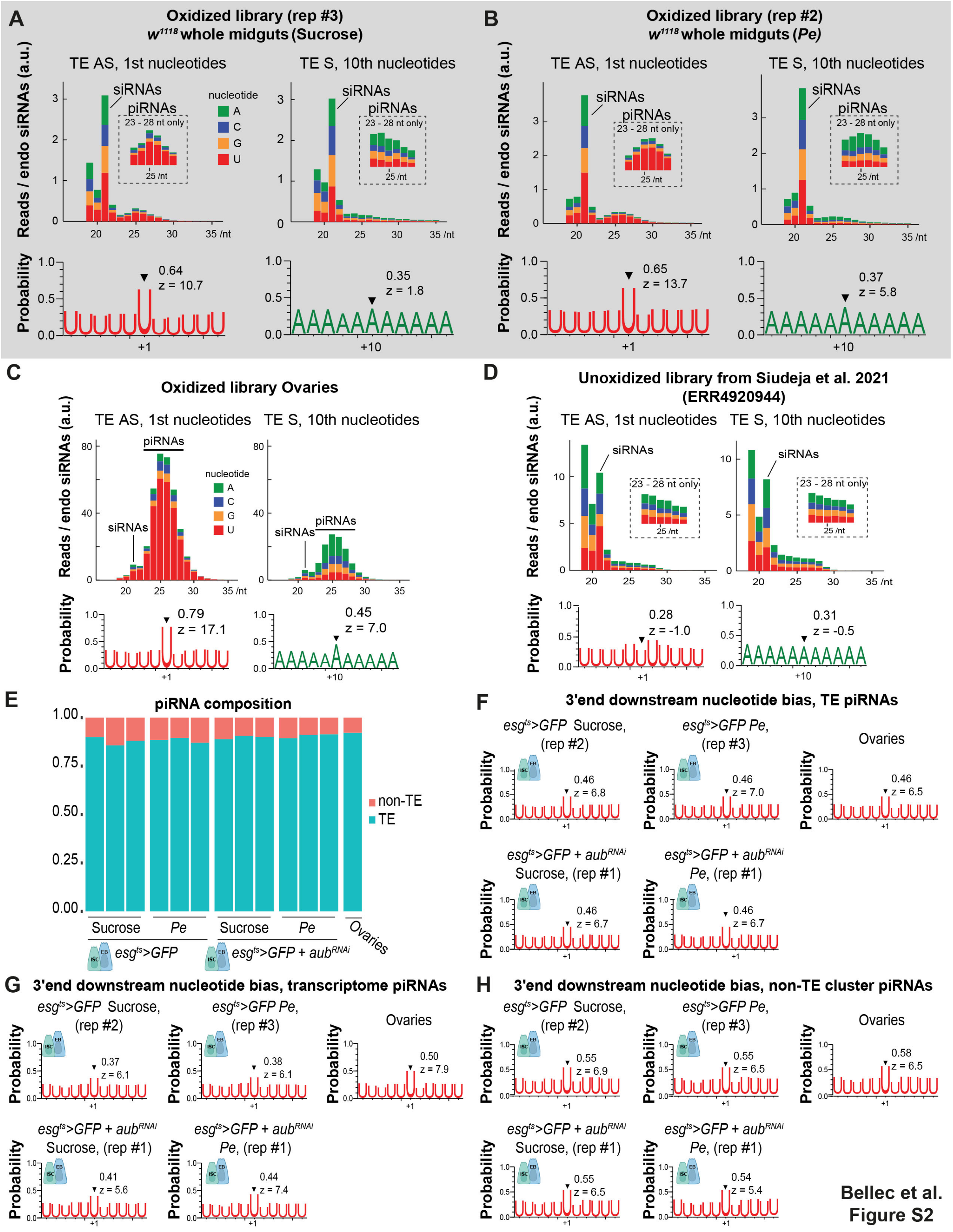


**Figure S2: Presence of piRNA-like signature in ISCs/EBs of the adult *Drosophila* midgut. Related to Figure 2. (A-D)** Size distribution (top graphs) of all transposon-mapping reads and the Uridine- and Adenine-frequencies of transposon-mapping reads of piRNA size (>22nt) (bottom graphs) in oxidized libraries from the whole intestine of control flies fed with sucrose (A) or *Pe* (B); in oxidized libraries from control ovaries (C) or in unoxidized libraries from the whole intestine as previously published (D).[68] siRNAs are typically sized at 21 or 22nt while piRNAs are longer than 22nt (see dashed black rectangles for higher magnification). Uridine (red) at the 1st base position of TE antisense piRNA reads (>22nt) is a common signature of all piRNAs while Adenine (green) at the 10th base position of TE sense piRNA reads is a signature of Ping-Pong piRNAs. **(E)** piRNA abundance in sorted ISCs/EBs. piRNAs mapping genomic TE sequences (as classified by RepeatMasker) or not (non-TE) are represented. **(F-H)** Uridine frequencies around the 3' end of transposon-mapping piRNAs (F); piRNAs derived from protein coding mRNAs (transcriptome) (G) or piRNAs originating from piRNA clusters excluding TE sequences. TE, transposable elements; AS, antisense; S, sense.

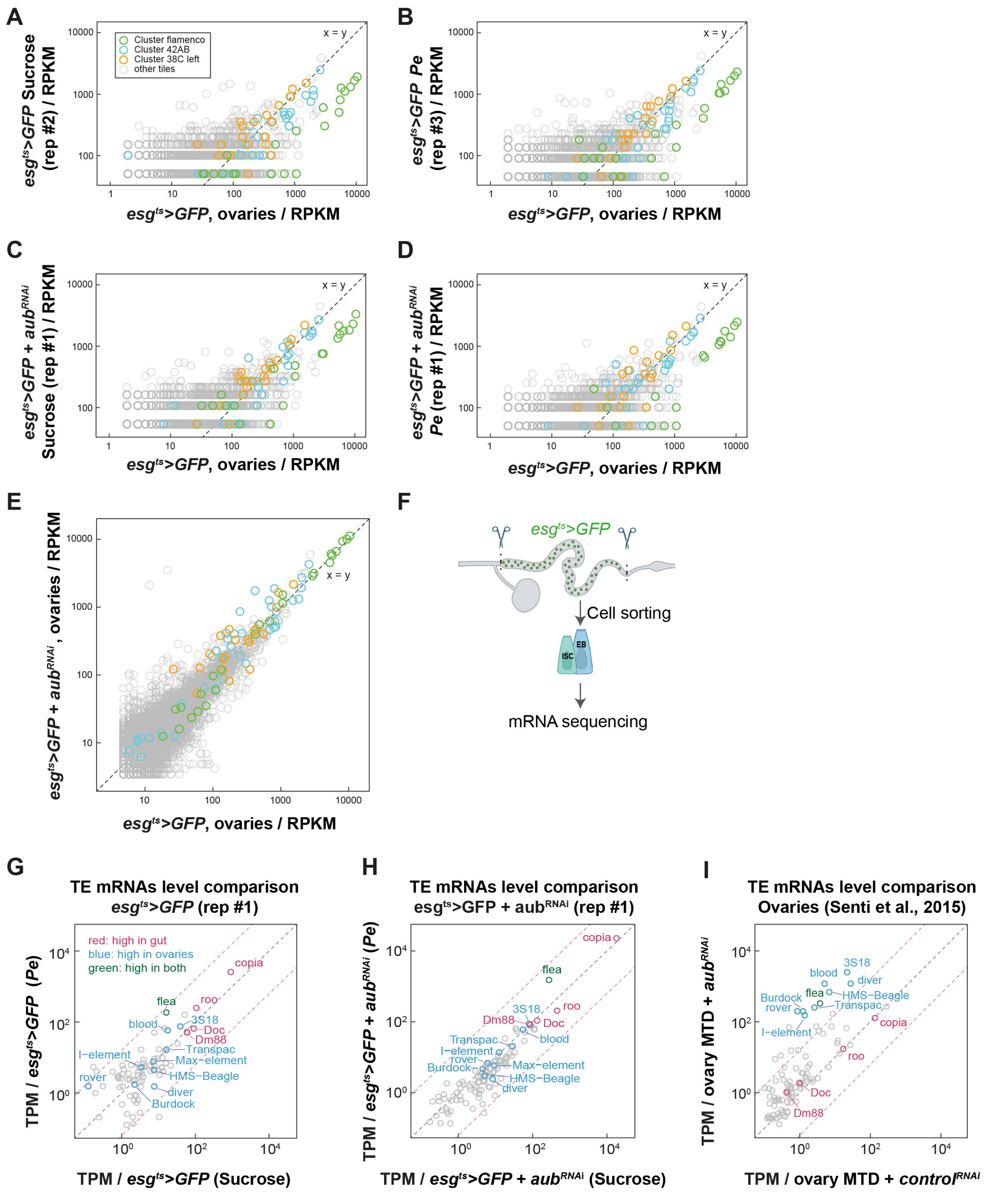


Bellec et al. Figure S3

**Figure S3: Transposable elements profile and piRNA-like signature in ISCs/EBs of the adult *Drosophila* midgut are distinct from that in ovaries. Related to Figure 2. (A-E)** Representation of the 0.5 kb tile analysis of piRNAs abundance measurement in sorted ISCs/EBs from control flies (A, B) or from flies expressing *aub*$^{RNAi}$ in ISCs/EBs (C, D) and fed with sucrose or *Pe,* compared to ovaries. The same analysis was performed to compare piRNAs abundance between ovaries from control flies and from flies expressing *aub*$^{RNAi}$ in ISCs/EBs (E). Tiles from piRNA clusters are coloured: cluster 38C in orange, cluster 42AB in blue and cluster flamenco in green. All other tiles are coloured in grey. **(F)** Schematic representation of the *Drosophila* intestine. The green dots represent ISCs/EBs that were sorted *via* FACs for mRNA sequencing. **(G-I)** Abundance of TE mRNAs in sorted control (G); or *aub*$^{RNAi}$ ISCs/EBs (H) from sucrose and *Pe* fed animals or from control and *aub*$^{RNAi}$ ovaries obtained from a previously published study (I).[69] TE mRNA counts are normalised to one million transcript counts including host mRNAs (TPM). Red dashed lines indicate 4-fold changes in either direction. Red, blue and green represent TE mRNAs highly expressed in the gut, in the ovaries or in both tissues, respectively. TE, transposable elements; AS, antisense; S, sense.

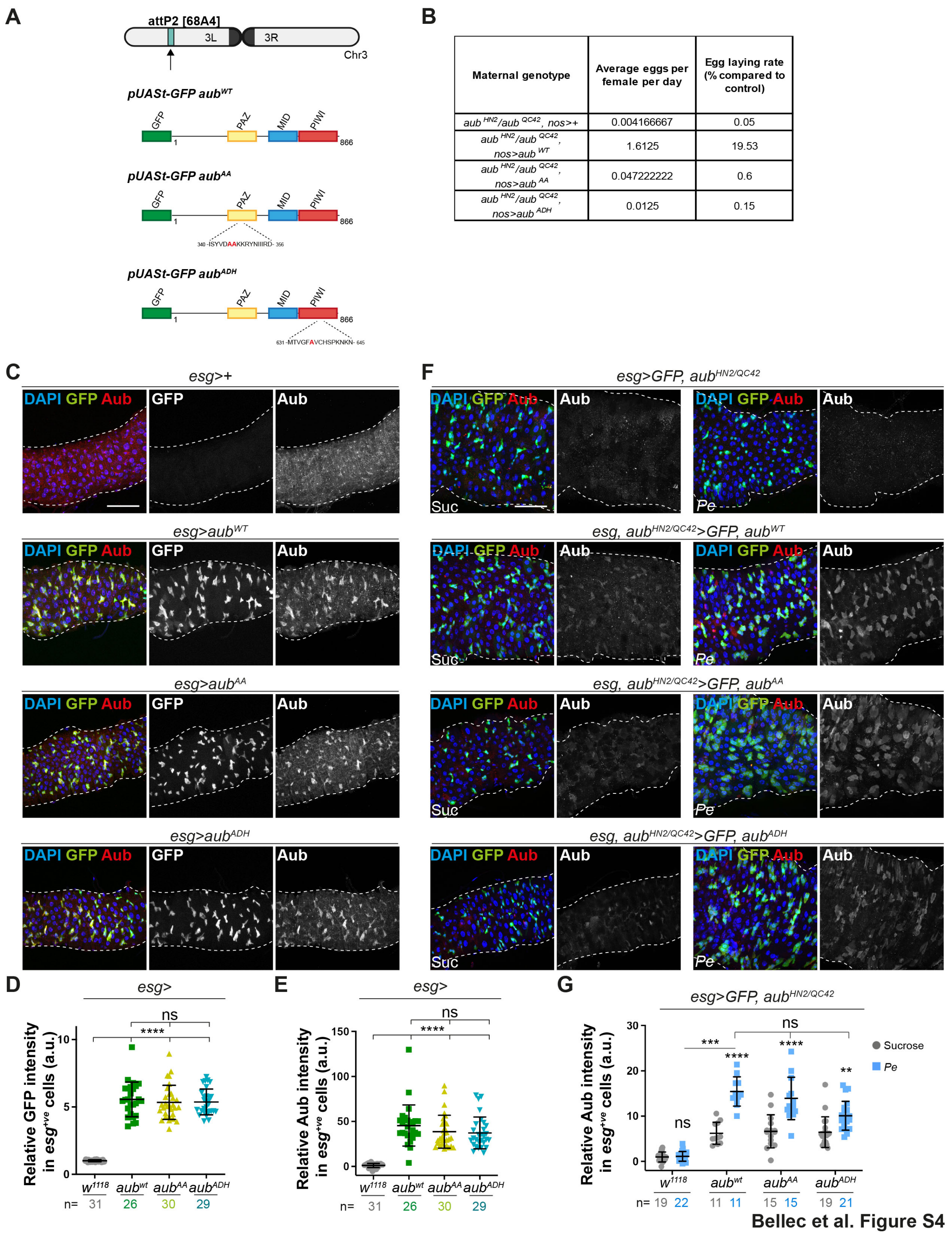


| Maternal genotype | Average eggs per female per day | Egg laying rate (% compared to control) |
|---|---|---|
| aub HN2/aub QC42, nos>+ | 0.004166667 | 0.05 |
| aub HN2/aub QC42, nos>aub WT | 1.6125 | 19.53 |
| aub HN2/aub QC42, nos>aub AA | 0.047222222 | 0.6 |
| aub HN2/aub QC42, nos>aub ADH | 0.0125 | 0.15 |

**Figure S4: Functional characterization of Aub transgenes in the adult midgut. Related to Figure 3. (A)** Schematic of the site-directed transgenesis induced to generate fly lines carrying inducible GFP tagged wild type Aub (*UAS-aub$^{WT}$*) or Aub mutant isoforms (*UAS-aub$^{AA}$* and *UAS-aub$^{ADH}$*). **(B)** Female fertility assays in *aub*$^{HN2}$/*aub*$^{QC42}$ mutant flies without or with expression of *aub*$^{WT}$, *aub*$^{AA}$ or *aub*$^{ADH}$ in the germline. **(C)** Aub staining (red) in posterior midguts from control flies or flies expressing *aub*$^{WT}$, *aub*$^{AA}$ or *aub*$^{ADH}$ within ISCs/EBs (green). Dashed white lines delineate the posterior midguts. **(D, E)** Quantification of GFP (D) or Aub staining (E) as in C. Shapiro-Wilk normality test followed by a Kruskal-Wallis one-way ANOVA and a Dunn's multiple comparisons test. n = number of midguts/flies quantified. **(F)** Aub staining (red) in posterior midguts of *aub*$^{HN2}$/*aub*$^{QC42}$ mutant flies expressing *aub*$^{WT}$, *aub*$^{AA}$ or *aub*$^{ADH}$ within ISCs/EBs. Dashed white lines delineate the posterior midguts. Nuclei are identified with DAPI. Scale bars =50µm. **(G)** Quantification of ISCs/EBs staining as in F. Unless otherwise noted, two-way ANOVA followed by Sidak's multiple comparisons tests were applied for statistical analysis. n = number of midguts/flies quantified. a.u., arbitrary units. Data are represented as mean +/- SD. ns, not significant; *$P < 0.05$, **$P < 0.01$, ***$P < 0.001$; ****$P < 0.0001$.

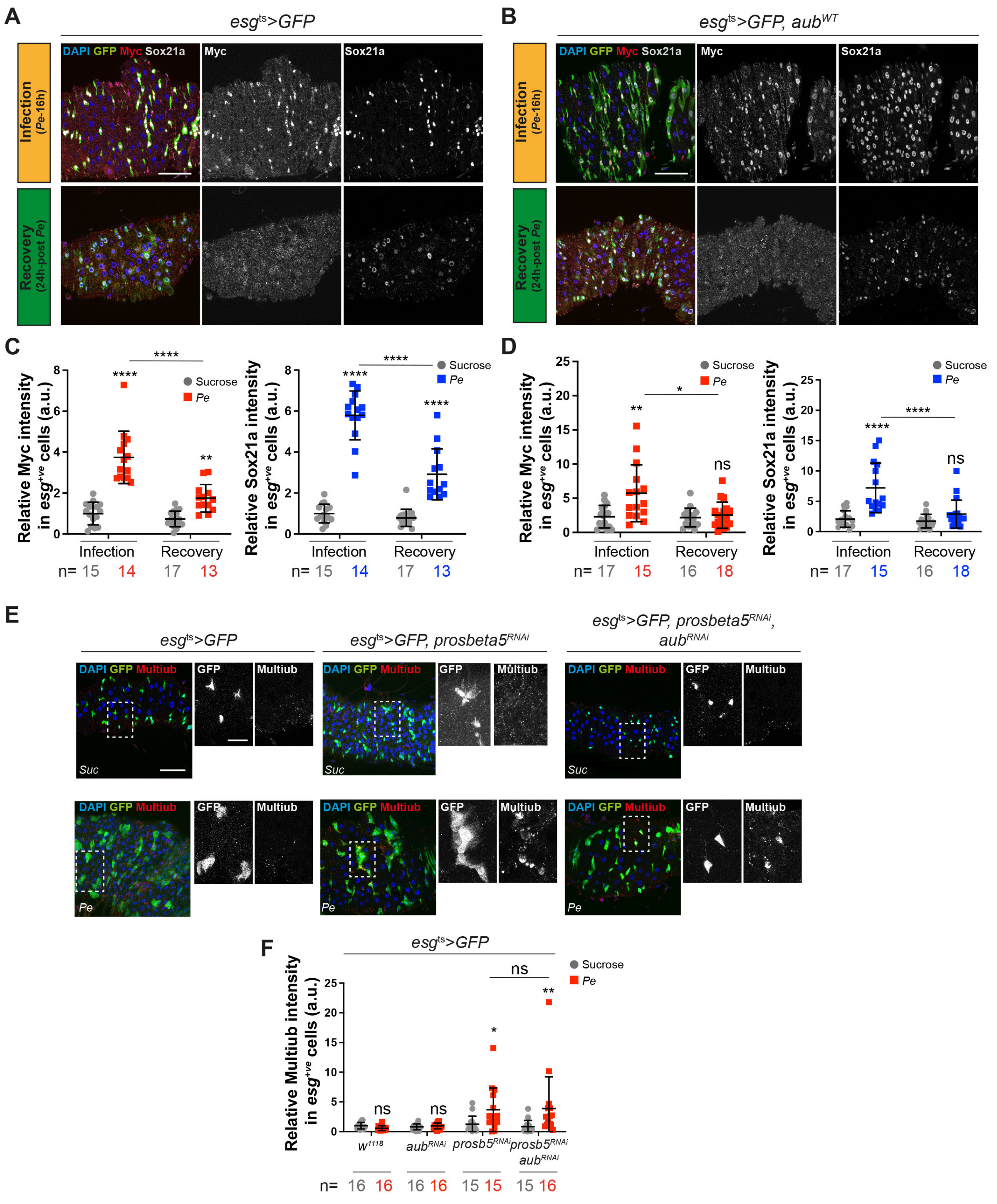


Bellec et al. Figure S5

**Figure S5: Aub does not impact protein stability in the regenerating midgut. Related to Figures 3 and 4. (A, B)** Myc and Sox21a staining in the posterior midguts of flies expressing GFP alone (A) or with *aub*$^{WT}$ (B) within ISCs/EBs (green) upon 16 hours of feeding with sucrose or *Pe* (upper panels) or upon 24 hours after removing Sucrose/*Pe* (bottom panels). **(C, D)** Quantification of Myc (red) and Sox21a (blue) within ISCs/EBs as in A and B, respectively. **(E)** Multiubiquitination staining (red) in posterior midguts of flies expressing GFP alone, or with *prosbeta5*$^{RNAi}$ or with *prosbeta5*$^{RNAi}$ and *aub*$^{RNAi}$ within ISCs/EBs (green) and fed with sucrose or *Pe*. Dashed white squares delineate the high magnifications shown in the right panels. Nuclei are identified with DAPI. Scale bars = 50μm. **(F)** Quantification of staining as in E. Two-way ANOVA followed by Sidak's multiple comparisons tests were applied for statistical analysis. n = number of midguts/flies quantified. a.u., arbitrary units. Data are represented as mean +/- SD. Data are represented as mean +/- SD. ns, not significant; $^*P < 0.05$, $^{**}P < 0.01$, $^{***}P < 0.001$; $^{****}P < 0.0001$.

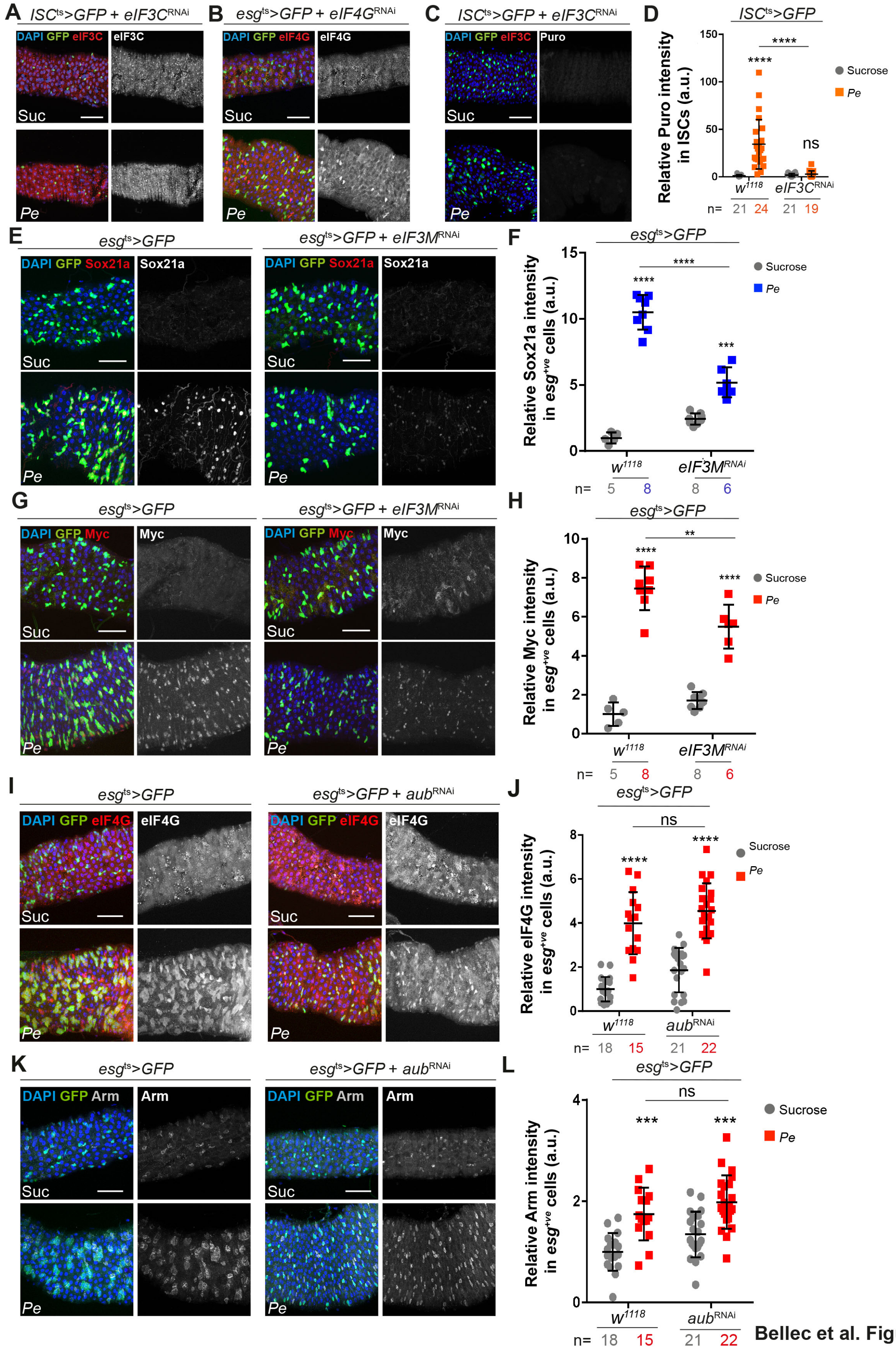


Bellec et al. Figure S6

**Figure S6: Selective function of Aub in the regulation of protein translation in ISCs. Related to Figure 5 and 6. (A, B)** eIF3C (A) and eIF4G (B) staining (red and grey) in posterior midguts of flies expressing GFP alone or with *eIF3C*$^{RNAi}$ (A) or *eIF4G*$^{RNAi}$ (B) within ISCs or ISCs/EBs, respectively, and fed with sucrose or *Pe*. **(C)** Puromycin staining (red and grey) in posterior midguts of flies expressing GFP alone or with *eIF3C*$^{RNAi}$ within ISCs (green) and fed with sucrose or *Pe*. **(D)** Quantification of staining as in C. **(E)** Sox21a staining (red and grey) in midguts expressing GFP alone or with *eIF3M*$^{RNAi}$ within ISCs/EBs (green) and fed with sucrose or *Pe*. **(F)** Quantification of staining as in E. **(G)** Myc staining (red and grey) in midguts expressing GFP alone or with *eIF3M*$^{RNAi}$ within ISCs/EBs (green) and fed with sucrose or *Pe*. **(H)** Quantification of staining as in G. **(I)** eIF4G staining (red and grey) in posterior midguts of flies expressing GFP alone or with *aub*$^{RNAi}$ within ISCs/EBs (green) and fed with sucrose or *Pe*. **(J)** Quantification of staining as in I. **(K)** Armadillo/β-Catenin staining (red and grey) in posterior midguts of flies expressing GFP alone or with *aub*$^{RNAi}$ within ISCs/EBs (green) and fed with sucrose or *Pe*. Nuclei are identified with DAPI. Scale bars= 50μm. **(L)** Quantification of Armadillo/β-Catenin staining as in K. Unless otherwise noted, two-way ANOVA followed by Sidak's multiple comparisons tests were applied. n = number of midguts/flies quantified. a.u., arbitrary units. Data are represented as mean +/- SD. ns, not significant; **P* < 0.05, ***P* < 0.01, ****P* < 0.001; *****P* < 0.0001.

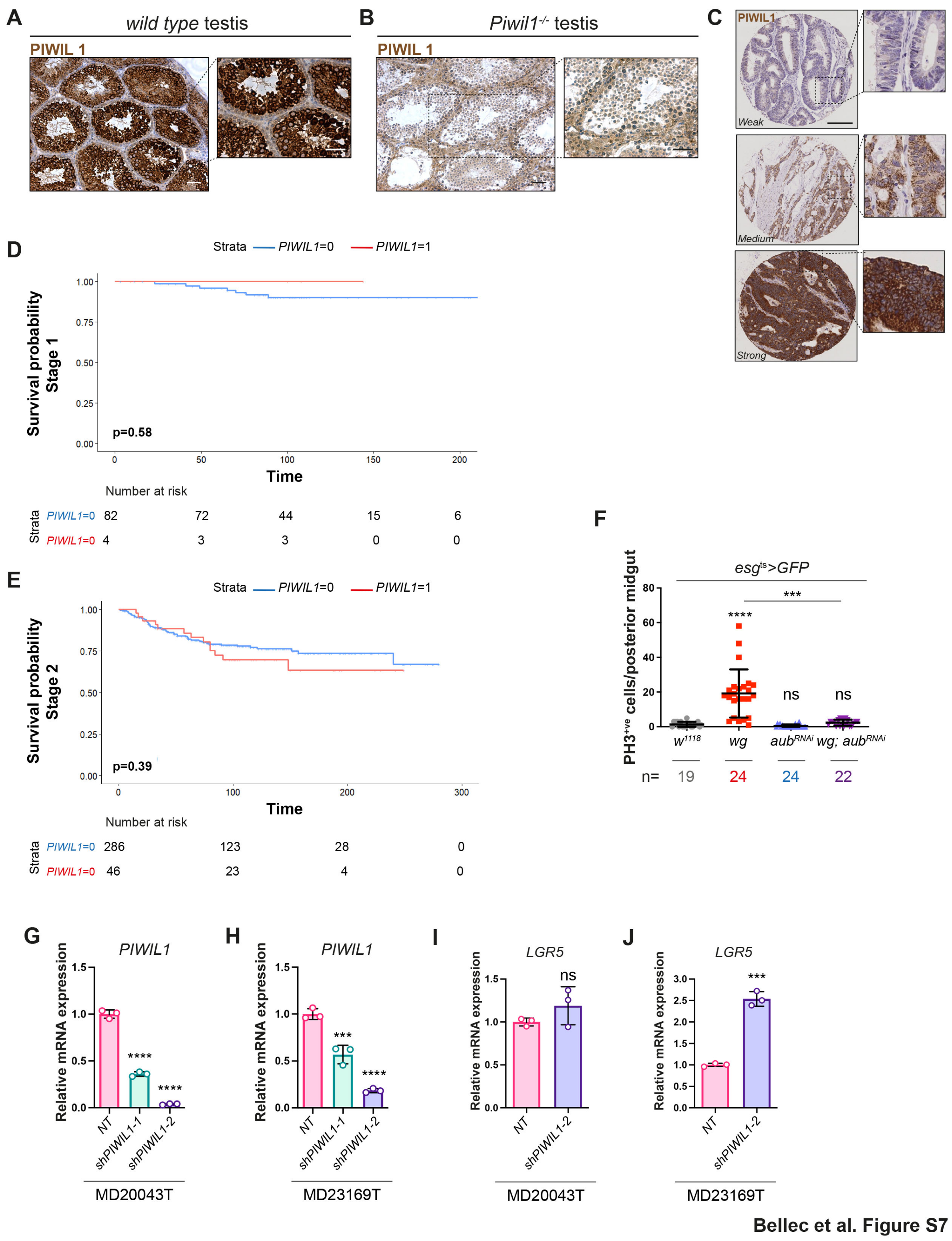


**Figure S7: *PIWIL1* expression in human CRC. Related to Figure 7.**

**(A, B)** PIWIL1 immunostaining in wild type and *PIWIL1* mutant testis. Dashed black squares delineate the high magnification shown in the right panels. Scale bars= 50µm. **(C)** Microscopic images of PIWIL1 expression in tissue samples from CRC patients.

Scale bar= 100 µm. **(D, E)** Kaplan-Meier survival analysis of *PIWIL1* expression in a cohort of CRC patients (n=787) showing the association between *PIWIL1* expression and cancer-specific survival in patients with stage 1 (D) and 2 (E) disease. **(F)** PH3-positive cells in the posterior midguts of control flies expressing GFP alone; with *Wg::HA* or $aub^{RNAi}$ or co-expressing *Wg::HA* and $aub^{RNAi}$ within ISCs/EBs. Shapiro-Wilk normality test followed by a Kruskal-Wallis one-way ANOVA and a Dunn's multiple comparisons tests were applied. n = number of midguts/flies quantified. **(G, H)** *PIWIL1* mRNA expression in intestinal organoids transduced with non-targeted control (NT), *shPIWIL1-1 or shPIWIL1-2* RNAi. One-way ANOVA and Dunnett's multiple comparisons tests. n = 3 biological replicates. **(I, J)** *LGR5* mRNA expression in intestinal organoids transduced with NT *or shPIWIL1-2* RNAi. Unpaired t test. n = 3 biological replicates. Data are represented as mean +/- SD. ns, not significant; ****P* < 0.001; *****P* < 0.0001.
